# Patient-derived lymphocytes drive smoldering lesion pathology in a chimeric multiple sclerosis mouse model

**DOI:** 10.1101/2025.09.16.676229

**Authors:** O. Perrot, M. Nordbeck, C. Bachelin, A. Luong, D. Roussel, D. Akbar, N. Sarrazin, A. Tenenhaus, C. Louapre, V. Zujovic

## Abstract

Multiple sclerosis (MS) is a chronic inflammatory disease of the central nervous system characterized by demyelination, axonal injury, and neurodegeneration. Smoldering lesions— defined by an inactive core, poor remyelination, and a rim of chronically activated microglia— are hallmarks of a progressive disease course and correlate with irreversible disability. The mechanisms driving their formation remain poorly understood.

Using a chimeric mouse model, we investigated the long-term impact of healthy donors (HD) and MS patient-derived lymphocytes (LY) graft on the evolution of spinal cord demyelinated lesion. Three months post-grafting, only MS-derived LY persisted as perivascular cuffs interacting with vascular and murine immune cells, mirroring MS smoldering lesion pathology. MS LY-grafted mice exhibited impaired functional recovery and slower somatosensory evoked potential (SSEP) conduction compared to controls. Electron microscopy confirmed glial scar formation, persistent demyelination and a lesion architecture with an inflammatory inactive center but active rim.

Single-nucleus RNA sequencing revealed an imbalance in CNS cell populations, with MS LY-grafted mice showing reduced neuronal abundance, enriched activated microglia, and immature oligodendroglial profiles, correlating with slower SSEP conduction speeds. Immune cell subclustering identified an enrichment of disease-associated microglia and interferon-responsive microglia in MS LY-grafted mice, marked by elevated pro-inflammatory markers and active myelin phagocytosis. Oligodendroglial cells displayed downregulated myelination and stress-response genes, with disrupted myelin ultrastructure confirmed by electron microscopy. Multiblock analysis revealed donor-specific variability, with some MS LY inducing outcomes akin to HD, while others drove severe pathology.

Our model enables mechanistic dissection of the transition from focal to diffuse CNS pathology in MS and captures patient-specific pathophysiological signatures, providing a platform for personalized mechanistic studies and targeted therapeutic strategies.

**Graphical abstract:** 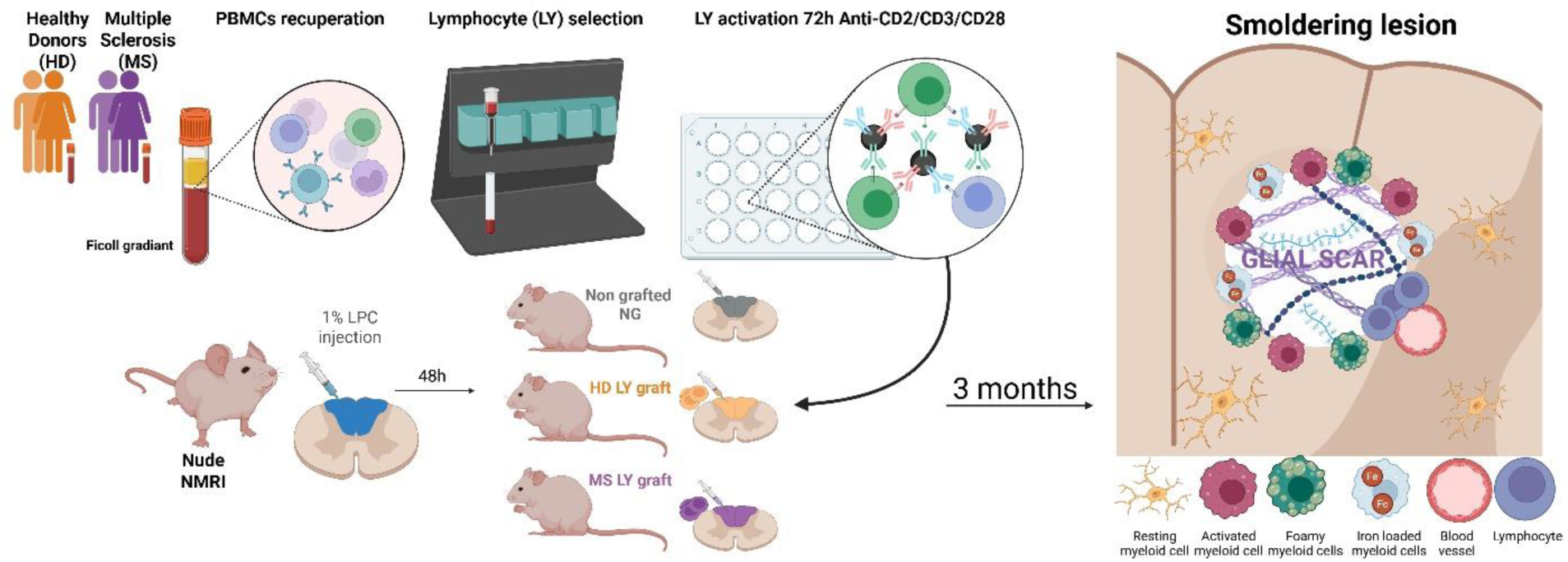

## Introduction

Multiple sclerosis (MS) is an autoimmune, neuro-inflammatory, and demyelinating disease of the central nervous system (CNS) (Compston & Coles, 2002) with different disease types. Relapsing/Remitting MS (RRMS) is the most common form (85%) and characterized by relapse-associated worsening (RAW) with periods of neurologic symptoms followed by partial or complete recovery, and possible progression independent of relapse activity (PIRA) (Woo et al., 2024). RRMS can evolve into a Secondary Progressive MS (SPMS) with accumulating disability without remission, including both RAW and PIRA. Primary Progressive MS (PPMS), which accounts for 15% of cases, features steady disability accumulation from disease onset and is primarily driven by PIRA.

Disease progression and symptomatology have been associated with different cellular and molecular events including waves of inflammatory demyelinating episodes followed by remyelination attempts that would lead to the formation of lesions. The study of MS *post-mortem* tissue led to the classification of different lesion types: (1) active lesions that present ongoing inflammation and demyelination, (2) inactive lesions that are demyelinated, with a glial scar but without active inflammation, and (3) “smoldering lesions” or mixed active/inactive lesions (Kuhlmann et al., 2017; Lucchinetti et al., 2000). These lesions are characterized by an inflammatory inactive and demyelinated center, glial scarring and an active rim compartment with ongoing inflammation, demyelination, and accumulation of either iron-loaded or myelin-loaded microglia (Absinta et al., 2021). Active lesions are more abundant in the early phases of RRMS and SPMS while smoldering lesions are more frequent in SPMS and PPMS (Frischer et al., 2015; Kuhlmann et al., 2017; Luchetti et al., 2018). The number of smoldering lesions is positively correlated with disability scores and lesion load at time of death, while being negatively correlated with the proportion of remyelinated lesions (Absinta et al., 2019; Luchetti et al., 2018). Ultimately, smoldering lesions represent a hallmark for patients with a severe disease course (Absinta et al., 2019).

Classically, experimental autoimmune encephalomyelitis (EAE) is used as an animal model of MS (Procaccini et al., 2015). It helped decipher the autoimmune component of MS and mimics the cyclical nature of RAW, characterized by episodes of neurological decline followed by periods of partial or complete recovery. This is crucial for understanding triggers and recovery processes. Other animal models improved our understanding of the different steps leading to remyelination, including the injection of various toxic agents into the CNS white matter tracts (e.g. lysolecithin, ethidium bromide and bacterial endotoxins), as well as cuprizone ingestion (Blakemore & Franklin, 2008). However, few models recapitulate continuous and irreversible progression of PIRA, which is essential for understanding how and why the disease shifts from a relapsing-remitting phase to a progressive one. Indeed, the initial infiltration of peripheral immune cells into the CNS sets off a chain reaction involving microglial cells that impedes remyelination and perpetuates inflammation and demyelination. This central activation of glial cells is crucial to the gradual expansion of smoldering-like lesions, making it essential to understand their impact on lesion development.

To address this challenging question, we established a new *in vivo* experimental paradigm to investigate how human MS lymphocytes (LY) influence remyelination and demyelination. We demonstrated that three weeks after grafting human LY into a demyelinated spinal cord lesion of nude mice, only MS patient derived LY hindered remyelination *in vivo*. By deciphering the mechanism of remyelination failure *in vitro*, we found that MS LY induced a higher pro-inflammatory activation of mouse microglial cells compared to healthy donor (HD) LY. This process leads to an early blocking of OPC differentiation. Moreover, we showed that patients LY influenced differentially the remyelination process, some showing a beneficial and some a deleterious effect on the repair process, a pattern mimicking the heterogeneity of remyelination capacity observed in MS patients (El Behi*., Sanson* et al., 2017).

In this study, we analyzed the long-term effects of the LY graft in a demyelinating lesion in the nude mouse spinal cord. Our results show that mice grafted with MS patients LY present mixed active-inactive lesions that share several aspects of smoldering-like lesions. This includes perivascular cuffs of human LY, inflammatory inactive and scared tissue at its center, failed remyelination, and accumulation of myelin- or iron loaded microglial cells at its border. The presence of these lesions in mice grafted with MS patients LY are correlated to behavioral deficits and slower axonal conduction, reflecting the chronic nature of the lesion. Interestingly, we also replicate a range of effects depending on the MS patient from whom the LY were isolated, mirroring the heterogeneity observed in patients.

## Material and methods

### Standard Protocol Approvals, Registrations, and Patient Consents

Approval for human blood sampling was obtained from the French Ministry of research (NCT03369106) and written informed consent was obtained from each study participant. All animal protocols are performed in accordance with the guidelines published in the National Institute of Health Guide for the Care and Use of Laboratory Animals, EU regulations (agreement n°C75-13-19) and the local “Charles Darwin” ethics committee.

### Patients and healthy donors

A total of 11 healthy donor (HD) and 21 MS patients paired by age and sex were included in the study (see table 1 and supplementary table 1 for details). All patients fulfill MS diagnostic criteria and individuals with any other inflammatory or neurological disorders were excluded from the study.

**Table 1:** Multiple sclerosis patients and healthy donor cohort description.

| Characteristic | MS patients | RRMS | PPMS | HD |
| --- | --- | --- | --- | --- |
| **n** | 17 | 16 | 1 | 7 |
| **Female/male** | 14/3 | 13/3 | 1/0 | 6/1 |
| **Mean age (range)** | 44 (23-70) | 43,63 (23-70) | 50 | 34,57 (25-51) |
| **[standard deviation]** | [12,99] | [13,32] |  | [8,17] |
| **Undertreatment/**<br>**treatment-free** | 7/10 | 7/9 | 0/1 |  |

*MS: Multiple sclerosis; RRMS: Relapsing/remitting multiple sclerosis; PPMS: Primary progressive multiple sclerosis; HD: Healthy donors. For details on treatment see supplementary table 1.*

### Collection and activation of patients and HD PBMCs and LY

80 ml venous blood was collected from MS patients or healthy individuals in acid citrate dextrose tubes. Peripheral blood mononuclear cells (PBMC) were extracted on a Ficoll gradient (GE Healthcare Life Sciences). Cells were washed in PBS (2×10 min at 1500 rpm) and culture medium (RPMI 1640, 10% fetal bovine serum (FBS); all products from ThermoFisher) (5 min at 1500 rpm). CD14 negative lymphocytes (LY) were selected by magnetic-activated cell sorting using anti-CD14 Microbeads (Miltenyi, 130-050-201) according to manufacturer instructions.

LY were plated at a density of 200,000 cells/24-well in 500 µl culture medium and activated using the human T cell Activation/Expansion kit (Miltenyi; 130-091-441), consisting in anti-CD2/anti-CD3/anti-CD28 bead-conjugated antibodies (at a density of 1 bead per 2 cells) in culture medium. After 72h of activation, cells were collected in RPMI medium at a concentration of 100,000 cells/µl for grafting.

### LPC injection and LY graft

Eight weeks old female Nude (RjOrl: NMRI-Foxn1^nu/^Foxn1^nu^) mice were purchased from Janvier (France). Focal demyelination was performed as previously described (El Behi*, Sanson* et al., 2017). Briefly, mice were anesthetized with a mix of ketamine (100 mg/ml; Imalgene 1000) and xylazine (10 mg/ml; Paxman). A demyelinating lesion was induced through stereotaxic injection of 1 µl, 1% lysophosphatidylcholine (LPC) (Sigma, 62962) in the dorsal horn of the spinal cord (between T9-T10). After 48h, 1 µl activated human LY (100,000 cells/µl) were grafted in the lesion site. Ten mice were grafted per individual donor. Ten mice underwent LPC injection without human lymphocyte graft and called the non-grafted group (NG).

### Behavioral testing

To assess the effect of the LY graft, we implemented two behavioral tests. Prior to LPC injection, the animals underwent a familiarization phase of three trials/day for five days. This pre-surgery readout was used as baseline performance to compare with the post-surgery results. The mice were assessed on the following tests at 1 week, 1 month, 2 months and 3 months after LPC injection.

The notched beam test allows us to assess mice motor coordination. The apparatus corresponds to a plastic beam that is one meter long and 8 mm wide, with a starting and ending plate of 80 mm and square slots (17 mm deep, 17 mm wide and 17 mm long, spaced every 17 mm and 30 cm high). For each trial, the number of errors and the time of crossing were recorded.

The grip test is a force measurement instrument to study neuromuscular function. The device allows the determination of the gram force of the front legs (using a bar) or the four legs (using a grid) of the mice. For each trial, the force in grams of the front legs and the four legs are recorded.

### Somatosensory evoked potential (SSEP)

Since the lesion affects the dorsal part of the spinal cord, where the ascending somatosensory fibers are located, somatosensory evoked potentials (SSEP) were recorded three months after LY graft, prior to tissue collection, to indirectly assess demyelination. The protocol was adapted from previously described approaches (Cloud et al., 2012; Farley et al., 2019). Recordings were performed under anesthesia with ketamine (100 mg/ml; Imalgene 1000) and xylazine (10 mg/ml; Paxman), with the animal secured on a pre-disinfected frame. Electrical stimulation was applied to the tibial nerve using disposable electrodes (Spesmedica, MN3512P150) placed on the hind paw (200 pulses, 0.1 ms duration, 3.5 Hz at 0-3 mA in 1 mA increments). Cortical responses were recorded via electrodes positioned over the somatosensory cortex and averaged across all stimuli. A ground electrode was placed subcutaneously above the cervical spinal cord of the mouse. Data acquisition and analysis were performed using Neuro-Mep.Net Omega software (Neurosoft).

We analyzed the SSEP latency as the onset of the response (Figure 3C left panel), and the mean was calculated across both hindlimbs. Conduction velocity was estimated by dividing the distance between stimulation and recording electrodes by the mean latency.

### Perfusion and tissue collection

Mice were anesthetized with a mix of ketamine (100 mg/ml; Imalgene 1000) and xylazine (10 mg/ml; Paxman). Euthanasia was performed after the assessment of SSEP with an overdose of pentobarbital (150 mg/kg; Euthasol).

For immunohistochemistry (IHC), mice were perfused intracardially with a solution of PBS 4% paraformaldehyde (PFA). Spinal cords were extracted and post-fixed for 1h. The tissue was cryoptrotected overnight in 20% sucrose solution, embedded in Cryomatrix (Epredia, 12868290), and frozen in cooled isopentane. The tissue was cryosectioned longitudinally to 12-µm sections and stored at −20 °C.

For electron microscopy (EM) mice were perfused with Phosphate buffer (PB) 4% PFA, 5% glutaraldehyde. Spinal cords were extracted and post-fixed for 1h. Spinal cords were cut into 1 mm sections and contrasted with 2% osmium tetroxide followed by 5% uranyl acetate before several steps of dehydration. Spinal cord sections were embedded in EponTM (Euromedex, 14120) and cut into semithin (0.5µm) and ultrathin (80nm) sections for analysis. Ultrathin sections were analyzed with a Hitachi HT7700 electron microscope.

When clearing the whole spinal cords, the tissue was post-fixed for 2-3h and stored in PBS with 0.1% sodium azide.

For single nuclei isolation, mice were euthanized by cervical dislocation after anesthesia and assessment of SSEP. The lesioned area of the spinal cord (1cm) was extracted and fresh-frozen on dry ice before storage at −80 °C.

### Immunohistochemistry and tissue clearing

Tissue sections were permeabilized and blocked for 1h at room temperature (RT) and stained with antibodies described in supplementary table 2. All primary antibodies were incubated overnight at 4°C, secondary antibodies were incubated for 1h at RT. Nuclei were counterstained with Hoechst (Sigma, 33342).

For whole tissue clearing, the tissue was dehydrated, bleached and rehydrated before staining. Tissues were incubated in primary antibody for 5 days at 37°C, washed multiple times a day and incubated with secondary antibodies for 5 days at 37°C. Tissues were cleared in Rapiclear solution (SunJin Lab, RC152001) overnight at 4°C. Nuclei were counterstained with Dapi dilactate (Sigma, D9564)

Images were acquired using a confocal microscope (Confocal A1R-HD25 Nikon). Microglia and oligodendrocytes were quantified in the perilesional rim, 3-months post-surgery. The lesion rim was defined by a 200 µm expansion of the lesion center.

### Nuclei isolation

For single nuclei RNA sequencing (snRNAseq), fresh frozen spinal cord tissue (∼1cm) was homogenized in 750 µl homogenization buffer (320 mM sucrose, 10 mM Tris pH 7.8, 5 mM CaCl_2_, 3 mM Mg(Ac)^2^, 1 mM β-mercaptoethanol, 0.1 mM EDTA, 0.1 mM PMSF, 0.1% NP40, 1% BSA, 1U/µl SUPERase inhibitor, in H_2_0) using a Dounce homogenizer on ice. Homogenization was performed with sequential strokes using a loose and tight pestle until the tissue was fully dissociated. The homogenate was filtered through 70 µm and 50 µm strainers, completed with homogenization buffer to a total volume of 2.7 ml, and subsequently mixed with 2.7 ml gradient buffer (50% OptiPrep, 10 mM Tris pH 7.8, 5 mM CaCl_2_, 3 mM Mg(Ac)^2^, 1 mM β-mercaptoethanol, 0.1 mM EDTA, 0.1 mM PMSF, 1% BSA, in H_2_0).

Nuclei were isolated on a sucrose gradient by gently layering the homogenate suspension on 5.4 ml sucrose cushion (29% OptiPrep, 71% cushion stock (250 mM sucrose, 150 mM KCl, 60 mM Tris pH 8, 30 mM MgCl_2_, in H_2_0), 1% BSA). Samples were subjected to ultracentrifugation at 7,000 rpm for 30 min using a SW41Ti rotor. Following centrifugation, the supernatant was aspirated, and nuclei pellets gently resuspended in recuperation buffer (1% BSA, 1U/µl SUPERase inhibitor in PBS).

Nuclei were passed through a 50 µm cell strainer and assessed for quality and concentration using trypan blue staining and an automated cell counter (Bio-Rad TC20). Only samples below 5% viability were selected for library preparation and sequencing.

### Single nuclei RNA sequencing

#### Library preparation and sequencing

Libraries were constructed using the GEM-X Universal 3′ Gene Expression v4 kit (10x Genomics) following the manufacturer’s instructions. Library quality and fragment size distribution were assessed using a TapeStation 4200 system (Agilent Technologies). Quantification of libraries was performed using a Qubit fluorometer (Thermo Scientific). Sequencing was carried out on a NovaSeq X Plus platform (Illumina). Eight libraries were loaded onto a single flow cell with a sequencing capacity of 10 billion reads. Paired-end sequencing was performed according to Illumina’s recommendations.

#### Pre-processing and alignment

FASTQ files were processed using the 10X Genomics CellRanger (v.9.0.0) (Zheng et al., 2017). Gene-barcode matrices were generated with the *count* function, including the -- include-introns flag and aligning to the GRCm39-2024-A reference mouse genome.

#### Ambient RNA correction

Ambient RNA contamination was corrected for each sample independently using CellBender (v.0.3.2) (Fleming et al., 2023). With the CellRanger *raw_feature_bc_matrix.h5* files as input, the *remove-background* function was used with half the learning rate (*learning-rate* 0.00005), while other parameters kept at default (Supplementary Figure 4).

#### Quality control and filtering

Nuclei were filtered uniformly across samples on a minimum detection of 300 features and 500 counts, and by excluding nuclei with > 7,000 features and > 30,000 counts. Nuclei with mitochondria content > 0.1 % and ribosomal content > 0.16 % were removed. Genes expressed in fewer than 5 nuclei were excluded. Doublets were identified based on co-expression of canonical marker genes in an initial clustering and subsequently removed (see below). After filtering, the dataset consisted of seven samples (one non-grafted but LPC-lesioned, two HD LY-grafted, and four MS LY-grafted) containing a total of 94k nuclei and 31k genes (Supplementary Figure 3).

#### Doublet removal, data processing, and clustering

The nuclei filtered and ambient RNA corrected count matrices were analyzed using the Seurat (v.5.3.0) (Hao et al., 2024) pipeline in R (v.4.3.3).

All samples were imported to generate a single Seurat object, with the “RNA” assay split by sample. Each sample was independently normalized and scaled using *NormalizeData()*, *FindVariableFeatures()*, and *ScaleData(vars.to.regress=“nCount_RNA”)* with default parameters.

Principal component analysis (PCA) was performed on variable features using *RunPCA()*, followed by sample integration using Harmony (v.1.2.3) (Korsunsky et al., 2019) via *IntegrateLayers(method = HarmonyIntegration, orig.reduction=“pca”, new.reduction=“harmony”)*. A uniform manifold approximation and projection (UMAP) was generated using *RunUMAP(dims=30, reduction=“harmony”)* for visualization.

To identify and remove potential doublets, an initial clustering analysis was performed on all nuclei to define broad cell types, followed by subclustering of each annotated cell fraction. Clusters were inspected for co-expression of canonical marker genes from distinct cell types, indicative of doublet events. Nuclei assigned to such mixed-marker clusters were removed and the dataset re-processed.

Broad cell-type clusters were identified using *FindNeighbors(dims=20, reduction=“harmony”)* and *FindClusters(resolution=2)*. Clusters were annotated based on canonical marker genes: neurons (*Rbfox3, Syn1, Grin2b*), astrocytes (*Slc1a2, Aqp4, Gfap*), vascular cells including endothelial cells, pericytes, and fibroblasts (*Col3a1, Flt1, Rgs5*), immune cells including microglia and peripheral mononuclear cells (*Ptprc, C1q, Cx3cr1*), OPC/COP (*Pdgfra, Cspg4, Vcan*), and oligodendrocytes (*Plp1, Mbp, Mog*).

To further resolve transcriptional states and subpopulations, the immune cell fraction and the OPC/COP plus oligodendrocyte fraction were re-integrated and clustered as described above. Immune cells were subclustered on the first 15 dimensions at a resolution of 1, while OPC/COP and oligodendrocytes were subclustered on 40 dimensions at a resolution of 0.3.

#### Cluster composition and correlation with lesion severity

To assess whether cluster abundance was associated with lesion severity, proportions were correlated with the speed of conduction through the spinal cord lesioned tissue using simple linear regression. Cell abundance was regressed against the speed of conduction, and the strength and significance of associations were reported using the coefficient of determination (R²) and corresponding p-values.

Data visualization and statistics was performed in R using the ggplot2 (v.3.5.2) and ggpmisc (v.0.6.1) package. For each cluster, scatter plots were generated with jitter applied to points to reduce overplotting, and regression lines with 95% confidence intervals overlaid (e.g. *geom_smooth(method = “lm”, se = TRUE)*; *stat_poly_eq(method=”lm”)*).

#### Marker gene identification

For marker gene identification, sample layers were joined via *JoinLayers()*. Markers were calculated using *FindAllMarkers(min.pct=0.25, logfc.threshold=0.5, only.pos=TRUE, test.use=”wilcox”)*. Only protein-coding genes were considered as markers.

#### Differential gene expression (DEG) analysis

For DEG analysis, sample layers were joined via *JoinLayers()*. DEG was calculated using MAST (v.1.32.0) (Finak et al., 2015) within the *FindMarker(min.pct=0.25, logfc.threshold=0.25, test.use=”MAST”)* function.

For each cell population, of interest, nuclei were subsetted according to cluster identity. Comparisons were made between experimental conditions (e.g. MS_slow vs. HD) or transcriptional states (e.g. DAM vs MG_hom). Genes with an absolute value of average log2 fold change ≥ 0.25 and a false discovery rate (FDR)-adjusted p < 0.05 were considered differentially expressed.

### Biostatistical analysis

The statistical analysis of snRNAseq data was performed in R according to the description above. Statistical tests regarding histological, physiological, or behavioral data were performed using Prism GraphPad software (GraphPad Prism version 10 for Windows). For behavioral data analysis, two-way ANOVA test was performed followed by a Tukey post-hoc test. For the analysis of histological data (EM and IHC) and SSEP, Kruskal-wallis test was performed. A Person test was performed for the correlation tests. The significance level was set at 0.05. To evaluate how individual donor lymphocytes (LY) influence graft outcomes, we performed a multiblock data analysis on behavioral, electrophysiological, and immunohistochemical variables. These variables were structured into two sets : a Behavioral and physiological block (Number of errors, Time to cross, % All limb strength, % Forelimb strength, Speed of conduction) and a biological block (Number of Mig+MOG+, % Mig+MOG+/Mig+, Number of Mig+FTL+, % Mig+FTL+/Mig+, Number of Mig+, Number of Olig2+, % Olig2+CC1+/Olig2+). To uncover shared information across these two blocks, we applied Regularized Generalized Canonical Correlation Analysis (RGCCA) (A. Tenenhaus et al., 2014, 2015; A. Tenenhaus & Tenenhaus, 2011; M. Tenenhaus et al., 2017) a statistical framework for data integration, implemented in the R package RGCCA (Girka et al., 2025). Within this framework, Multiple Co-Inertia Analysis (MCOA) (Chessel & Hanafi, 1996) can be specified as one possible configuration. A key advantage of MCOA is its ability to represent individuals in a consensus space that integrates all modalities, thereby facilitating both visualization and interpretation. Here, we considered MCOA with two components. To assess the reliability of the weights which reflect the contribution of each variable to the consensus space, we applied a bootstrap procedure. Specifically, bootstrap samples of the same size as the original dataset were repeatedly drawn with replacement, and MCOA was applied to each. From these estimates, we calculated standard deviations, bootstrap confidence intervals, t-ratios (the ratio of the parameter estimate to the bootstrap-estimated standard deviation), and p-values (assuming the t-ratio follows a standard normal distribution). Because multiple p-values were tested simultaneously, false discovery rate (FDR) correction was applied, and significant adjusted p-values were visualized with stars.

## Results

### MS patient’s LY form perivascular cuffs

We clarified the spinal cord of grafted mice to detect the presence or absence of human LY (HLA, red) and their localization in the vicinity of blood vessels (Collagen IV, white) or mouse myeloid cells (mix F4/80; CD11b; CD68, green) (Figure 1A-E). While we did not detect human LY in HD grafted mice (Figure 1A; C), we detected human MS patient LY in the perivascular area (Figure 1B; D; E). Interestingly, a 3D reconstruction enabled us to not only observe these LY perivascular cuffs but an accumulation of murine myeloid cells in close contact with these human cells (Figure 1D; E; mix F4/80; CD11b; CD68, green, arrows). The presence of MS patients LY around blood vessels was further confirmed by EM (Figure 1F-G). These observations confirm that after 3 months MS patients LY interact with mouse immune cells and might be implicated in the persistence of inflammation and a deleterious environment impeding the remyelination process.

**Figure 1:**
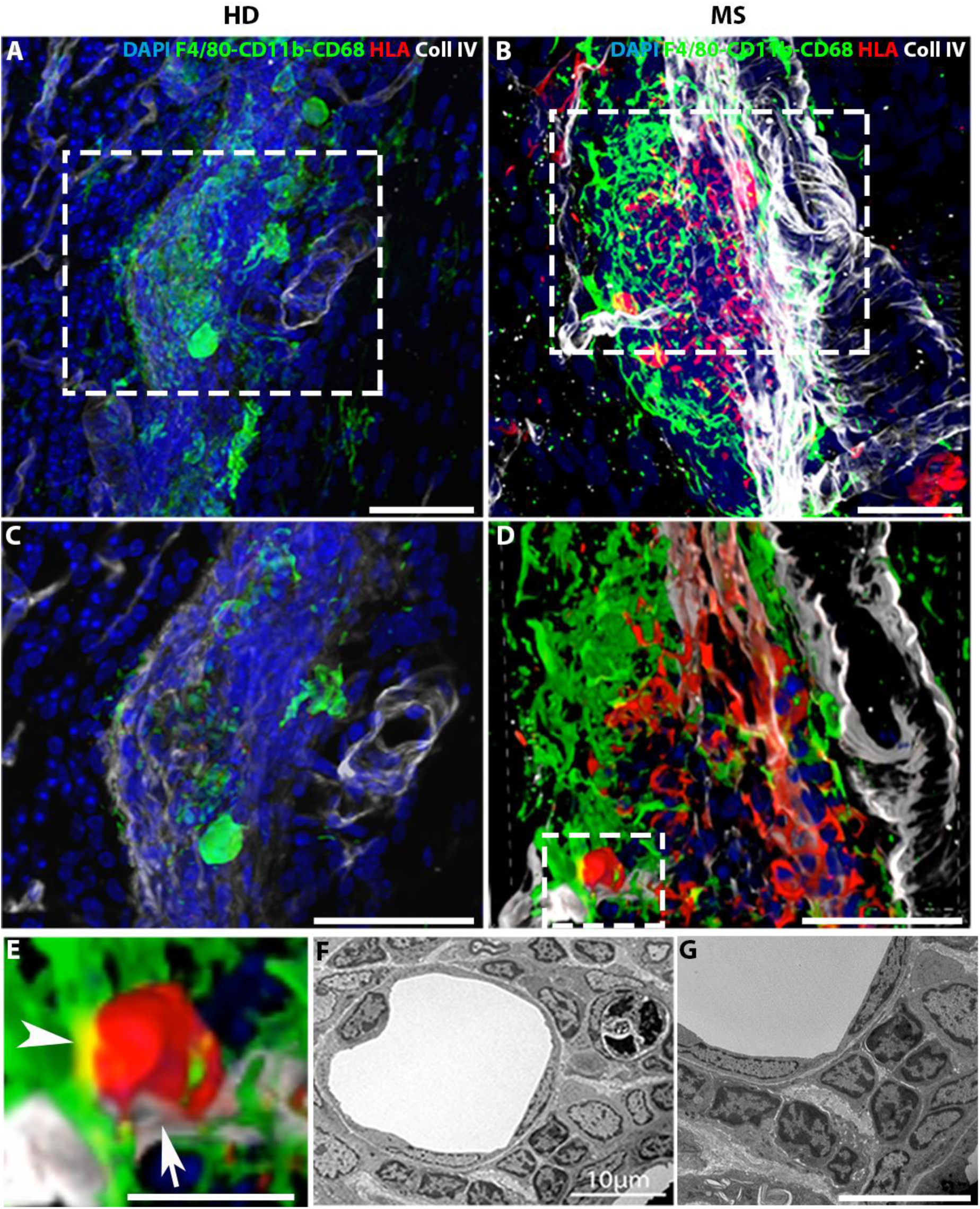
Human LY from MS patients, still present 3 months after the graft, form perivascular cuff and are in close contact with mouse myeloid cells. **(A-E)** Three-dimensional confocal image of mice grafted with (A; C) HD LY or mice grafted with (B; D; E) MS LY. Spinal cords were cleared using a modified iDISCO/RapiClear protocol. Scale bar: 50µm. (A) 3D image of HD grafted mice, (C) Zoom in the 3D confocal images. We did not observe LY in HD grafted mice, and only a few mouse myeloid cells (green). (B) 3D image of MS grafted mice, (D; E) Zoom in the 3D confocal images. We observed MS LY (red) in close contact to mouse myeloid cells (green, arrowhead) and around blood vessels (white, arrow), Scale bar: 20µm (A-D) or 20µm (E). **(F-G)** Semi thin section of a mouse grafted with MS LY. We observed LY around blood vessels. Scale bar: 10µm.

### MS patient’s LY induce scar formation and impede remyelination

Since our previous results, 3 weeks post-surgery, had already revealed perturbed remyelination in mice grafted with MS patient LY (El Behi*., Sanson* et al., 2017), we next examined the evolution of remyelinated and scarred areas by electron microscopy at 3 weeks and 3 months post-surgery in NG, HD, and MS grafted mice (Supplementary Figure 1A-I,). In the NG and HD groups, no significant changes were observed over time. In contrast, MS grafted mice showed a significant decrease in the remyelinated area (Supplementary Figure 1J) and an increase in glial scar area (Supplementary Figure 1K). When comparing different mice at 3 weeks and 3 months grafted with LY from the same individual, we observed a decrease in the remyelinated area (Supplementary Figure 1M) and an increase of glial scar area (Supplementary Figure 1N). Along with a decrease of macrophages over time (Supplementary Figure 1G; H; I, macrophages in blue; Supplementary Figure 1L), our results indicate a progression from peripheral attack to CNS dysregulation.

Next, we focused on the experimental time point 3 months post-surgery in NG mice (Figure 2A, D, G, J), mice grafted with HD LY (Figure 2B, E, H, K) or with MS LY (Figure 2C, F, I, L). We observed that remyelination was completed in NG mice (Figure 2A, D) and HD LY grafted mice (Figure 2B, E) as illustrated by the presence of axons remyelinated either by oligodendrocytes (OL) (Figure 2J, K, green) or Schwann cells (Figure 2 J, K, blue). In contrast, MS patient LY-grafted mice displayed sustained demyelination with the formation of a glial scar at the center of the lesion (Figure 2C, F, I). While there were some remyelinated axons at the lesion’s border (Figure 2I, L, blue, by Schwann cells), most of the axons remained unmyelinated (Figure 2I, L, yellow). Quantifications of the remyelinated and scar areas confirmed a lower percentage of remyelinated area in MS LY-grafted mice compared to NG or HD LY-grafted mice (Figure 2M). MS LY-grafted mice had a larger glial scar area compared to NG or HD LY-grafted mice (Figure 2N). These data demonstrate that MS patient LY induced the formation of a glial scar and impeded the remyelination process.

**Figure 2:**
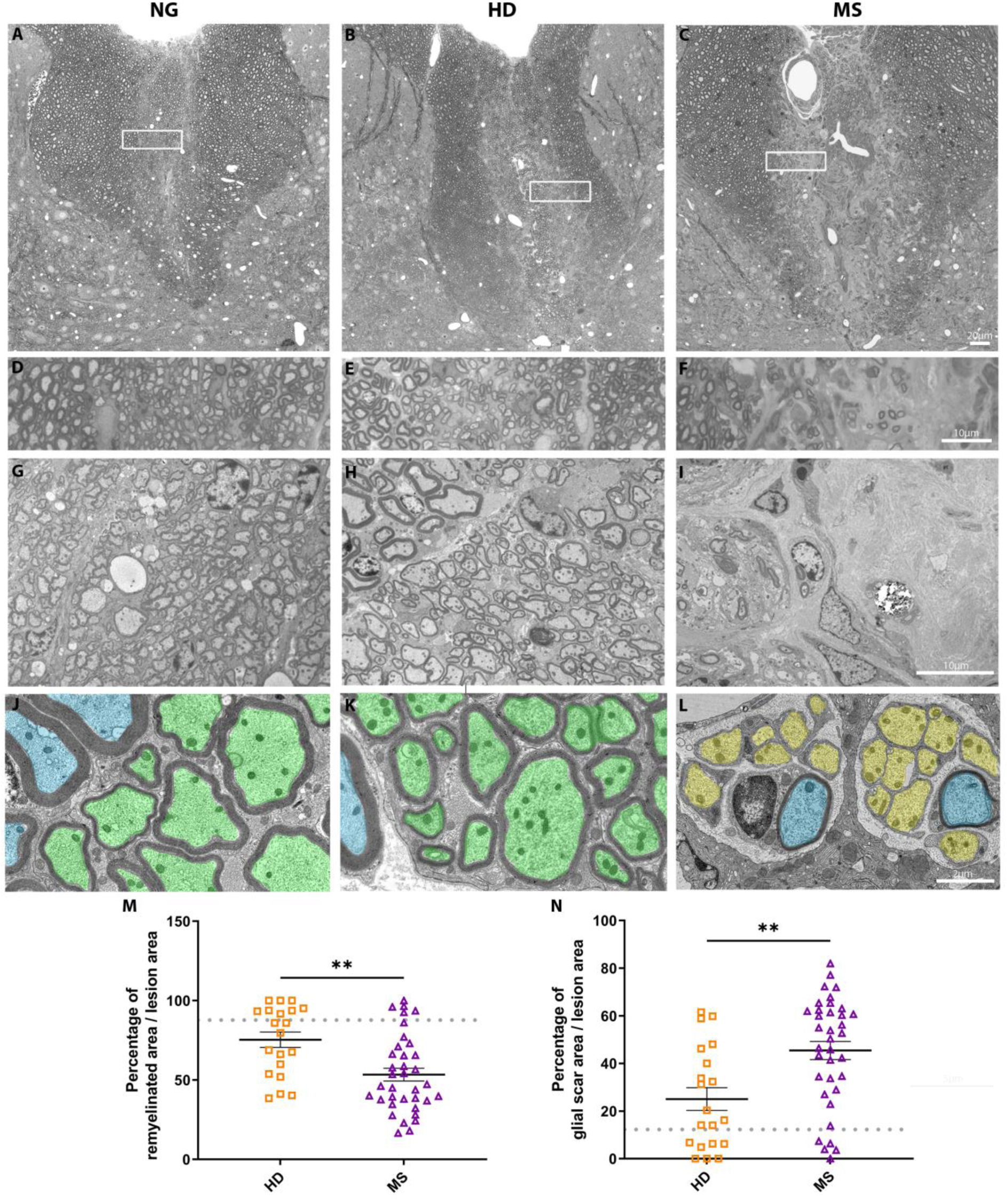
MS patient LY graft induces a chronic demyelinated lesion with scaring tissue. Electron microscopy of NG (A, D, G, J; n=6 mice, gray), HD LY grafted (B, E, H, K; n= 15 mice, 6 donors, orange), and MS LY grafted mice (C, F, I, L; n= 35 mice, 14 donors, purple), 3-month post-surgery. **(A-C)** Semi thin section of the dorsal horn of the spinal cord lesion. **(D-F)** Zoom inside the lesion. **(G-I)** Zoom inside the lesion center highlighting remyelinated axons in NG and HD LY grafted mice while MS LY grafted mice developed a glial scar. **(J-L)** Ultrathin sections of the lesion border highlighting remyelinated axons by oligodendrocytes (green) and Schwann cells (blue) in NG and HD LY grafted mice or unmyelinated axons (yellow) in MS LY grafted mice **(M)** Mean percentage of remyelinated area in reference to total lesion area. **(N)** Mean percentage of glial scar area in reference to total lesion area. Each dot represents one mouse; the mean and SEM are overlaid in black. The dotted gray line represents the mean value of the NG group. One way ANOVA followed by Tukey’s post-hoc: **: p<0,01.

### Longitudinal behavioral analysis reveals impaired recovery in mice grafted with MS patient LY

We monitored lesion evolution through a longitudinal assessment of sensory-motor function using behavioral tests in the 3 groups of mice (Supplementary Figure 2A, NG mice gray, HD LY grafted mice orange, MS patient LY grafted mice purple). Baseline performance was recorded one week before surgery, followed by testing at 1 week, 1 month, 2 months, and 3 months post-surgery.

For the notched beam test, there were no significant differences between the duration of crossing the beam at any time point (Supplementary Figure 2B). However, 1-week post-surgery, all groups made significantly more errors compared to the baseline assessment (Figure 3A). NG and HD LY-grafted mice subsequently recovered and even outperformed their baseline levels at 3 months post-surgery. MS LY-grafted mice continued to make significantly more errors compared to NG, HD, or their own baseline values at 3 months post-surgery.

**Figure 3:**
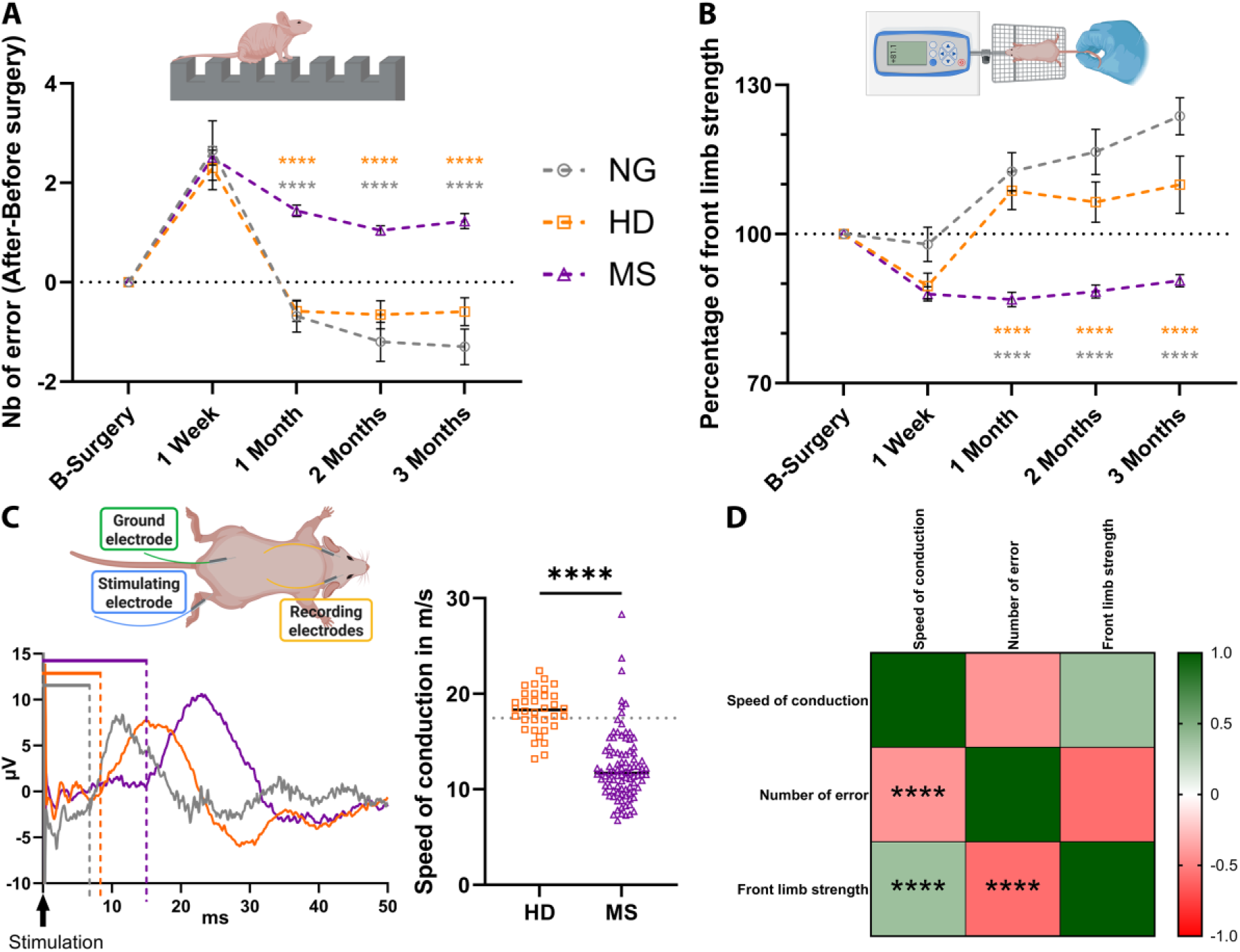
Mice grafted with MS LY show impaired behavioral and physiological performances. **(A-B)** Behavioral testing was assessed for NG mice (n=20 mice, gray), HD LY grafted (n= 27 mice, 3 donors, orange) and MS LY grafted mice (n= 77 mice, 9 donors, purple). For each time point, the MS group was compared to the NG (gray stars) or HD (orange stars) group. 2-way ANOVA followed by Tukey’s post-hoc. (A) Notched beam test and graph representing the mean number of errors and SEM post-surgery in reference to baseline performance. (B) Grip test and graphs representing relative front limb strength compared to baseline performance. **(C)** Left panel: Experimental set up of measuring SSEP conduction velocity. Briefly, an electrical stimulation is performed on the tibial nerve while the response to the stimuli is measured near the somatosensory cortex area. The arrow represents the artifact of stimulation. Dotted lines represent the beginning of the cortical response. Right panel: Mean conduction speed in m/s for each of the HD (n= 34 mice, 6 donors, orange) and MS group (n= 94 mice, 17 donors, purple). The gray dotted line represents the mean conduction velocity of NG mice (n= 20 mice) One way ANOVA followed by Tukey’s post-hoc. **(D)** Correlation table using the Pearson test of correlation. Significance is represented as stars: *: p<0,05; **: p<0,01; ***: p<0,001; ****: p<0,0001.

In the grip strength test, no differences in total limb strength were detected across experimental conditions (Supplementary Figure 2C). Nevertheless, MS LY-grafted mice displayed persistently reduced forelimb strength at 1-, 2-, and 3-months post-surgery compared to NG or HD LY-grafted mice (Figure 3B). NG and HD LY-grafted mice regained and improved strength over time, whereas MS LY-grafted mice failed to recover. These results indicate that MS LY-grafted mice did not functionally recover after lesion induction.

Given that the demyelinating lesion is induced in the dorsal horn of the spinal cord where sensory fibers are located, we measured the somatosensory evoked potential (SSEP) by recording the latency of the somatosensory cortex response to a hind paw stimulation (Figure 3C). The SSEP represents therefore an indirect evaluation of the lesion load in the somatosensory tract. Representative traces show a delayed response in MS LY-grafted mice (Figure 3C left panel). This delay was confirmed across all mice, as shown in the graph summarizing the mean conduction speeds for the three groups with a significant difference compared to NG mice or HD LY grafted mice (Figure 3C right panel). Correlation analysis revealed that the conduction speed positively correlated with the forelimb strength (r=0.38; p<0.01) and negatively correlated with the number of errors made on the notched beam (r=-0.41; p<0.01) (Figure 3D). Thus, slower conduction speed was associated with poorer behavioral performance.

We next examined the effect of human LY at the donor level by comparing the speed of conduction between mice grafted with LY from the same individual (Supplementary Figure 2D). Results revealed donor-specific effects, reflecting heterogeneity in remyelination capacity: some MS donors induced outcomes comparable to healthy donors, while others caused remyelination failure and a persistently slow conduction velocity. Accordingly, we classified MS donors as “slow” (SSEP speed of conduction < 12 m/s) or “fast” (12 - 17 m/s) when the conduction speed was closer to that of HD LY-grafted mice (> 17 m/s). The average speed of conduction for NG mice was 16.4 m/s while non-lesioned baseline velocity was measured at 19.75 m/s.

### Imbalance of cell populations in MS LY grafted mice

To assess how human LY grafting impacted mouse CNS environment, we performed single-nucleus RNA sequencing (snRNAseq) on lesioned/grafted spinal cords. We analyzed 2 mice grafted with LY from 2 different HD donors, and 4 mice grafted with LY from 4 different MS patients. The MS donors were selected based on SSEP conduction speed, to represent distinct functional states: 2 donors with slow SSEP conduction (MS_slow) and 2 with fast SSEP conduction (MS_fast) (Supplementary Figure 2D). This allowed us to compare graft effects across contrasting electrophysiological profiles.

Unsupervised clustering analysis led to the identification of all major CNS cell types (Figure 4A). To investigate disease-related alterations, we compared MS and HD samples at both cellular abundance and transcriptional levels, enabling the identification of differentially expressed genes and dysregulated pathways associated with disease states.

**Figure 4:**
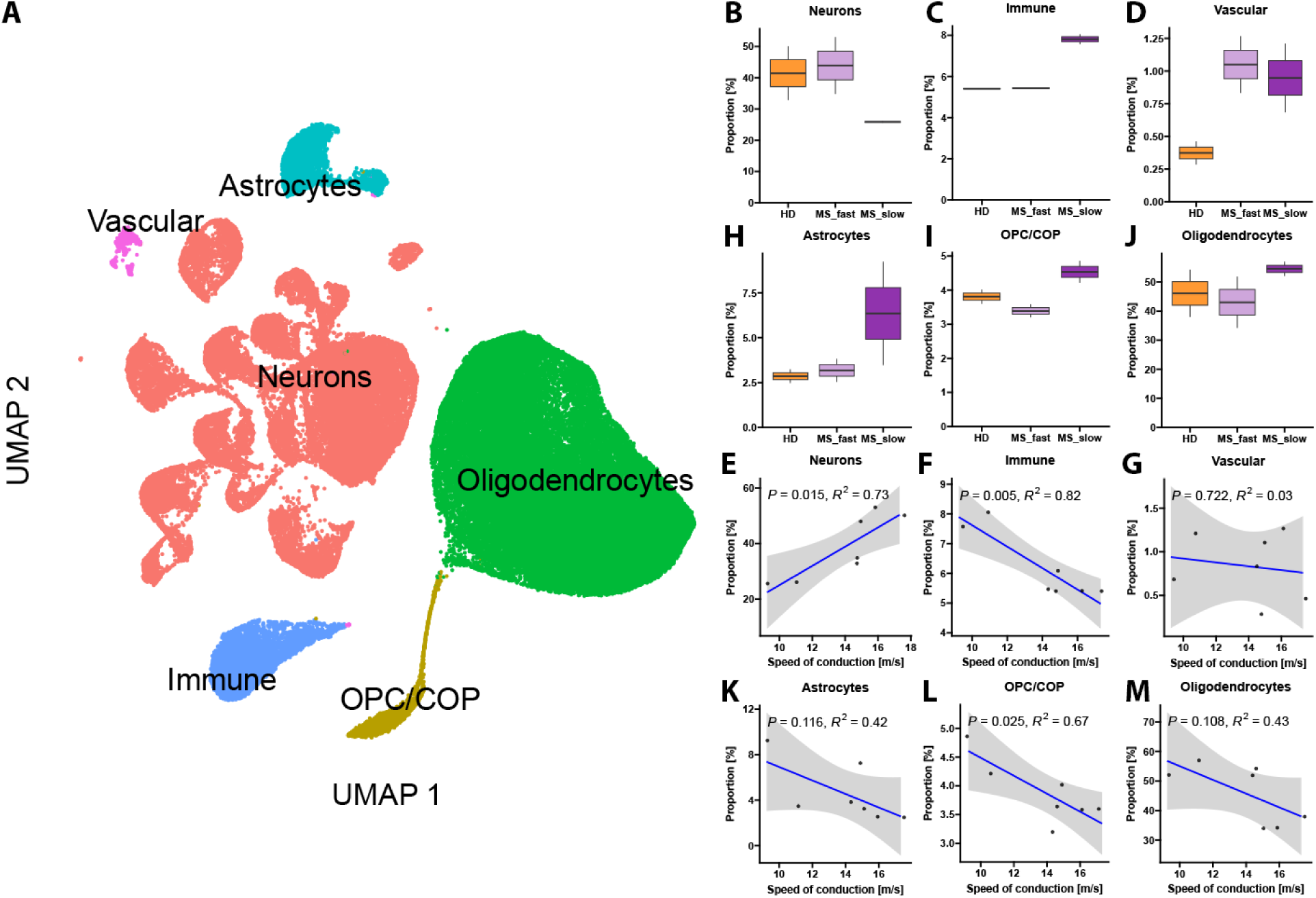
Imbalance of cell populations in MS LY grafted mice. **(A)** UMAP of major CNS populations after clustering analysis (n = 7 samples,2 HD, 4 MS (2 samples associated to fast and 2 to slow SSEP conduction velocities), 3 months post-surgery, 94,006 nuclei). **(B-D; H-J)** Boxplots highlight relative cell abundance of (B) neurons, (C) immune cells, (D) vascular cells, (H) astrocytes, (I) OPC/COP, and (J) oligodendrocytes. HD samples are shown in orange, MS samples associated with fast SSEP speed of conduction in light purple, and MS samples associated with slow SSEP speed of conduction in dark purple. **(E-G; K-M)** Cell abundance regressed against the speed of conduction. Strength and significance of associations were reported using the coefficient of determination (R²) and corresponding p-values.

Among the MS groups, MS-slow mice showed the greatest divergence from HD, represented by a reduction in neuronal abundance and an increase in immune cells, astrocytes, OPC/COP, oligodendrocytes, and vascular cells relative to both HD and MS-fast mice (Figure 4B–D; H-J). Interestingly, the SSEP assessed speed of conduction correlated positively with neuronal abundance and negatively with the abundance of immune cells, astrocytes, and OPC/COP (Figure 4E-G; K-M).

### Emergence of disease associated microglia at the lesion rim after MS LY graft

Subclustering of immune cells revealed distinct populations of microglia, monocytes/macrophages, lymphocytes, and neutrophils, defined by marker gene expression (Figure 5A-B, Supplementary Figure 5). Microglia were globally overrepresented in MS_slow subgroup, with enrichment of homeostatic, disease-associated (DAM), and interferon-responsive (IRM) subtypes (Figure 5C-E). Among these, only DAM abundance correlated negatively with the speed of conduction (Figure 5F-H, Supplementary Figure 6). To assess their potential impact in the lesion environment, we compared DEGs between homeostatic microglia and the DAM or IRM subtypes. DAM were characterized by upregulation of ApoE, Spp1, Ftl1, Gpnmb, chemokines (e.g., Ccl3, Ccl4), and Pdgfa, alongside reduced expression of canonical homeostatic markers (P2ry12, Ccr5, Cx3xr1, Tmem119) (Figure 5I). IRM displayed elevated interferon-signaling genes (Irf5, Irf7, Ifi202b, Ifit2, Ifit3, Ifi204, OAS family genes), chemokines Ccl2 and Ccl4, and reduced P2ry12 expression (Figure 5J).

**Figure 5:**
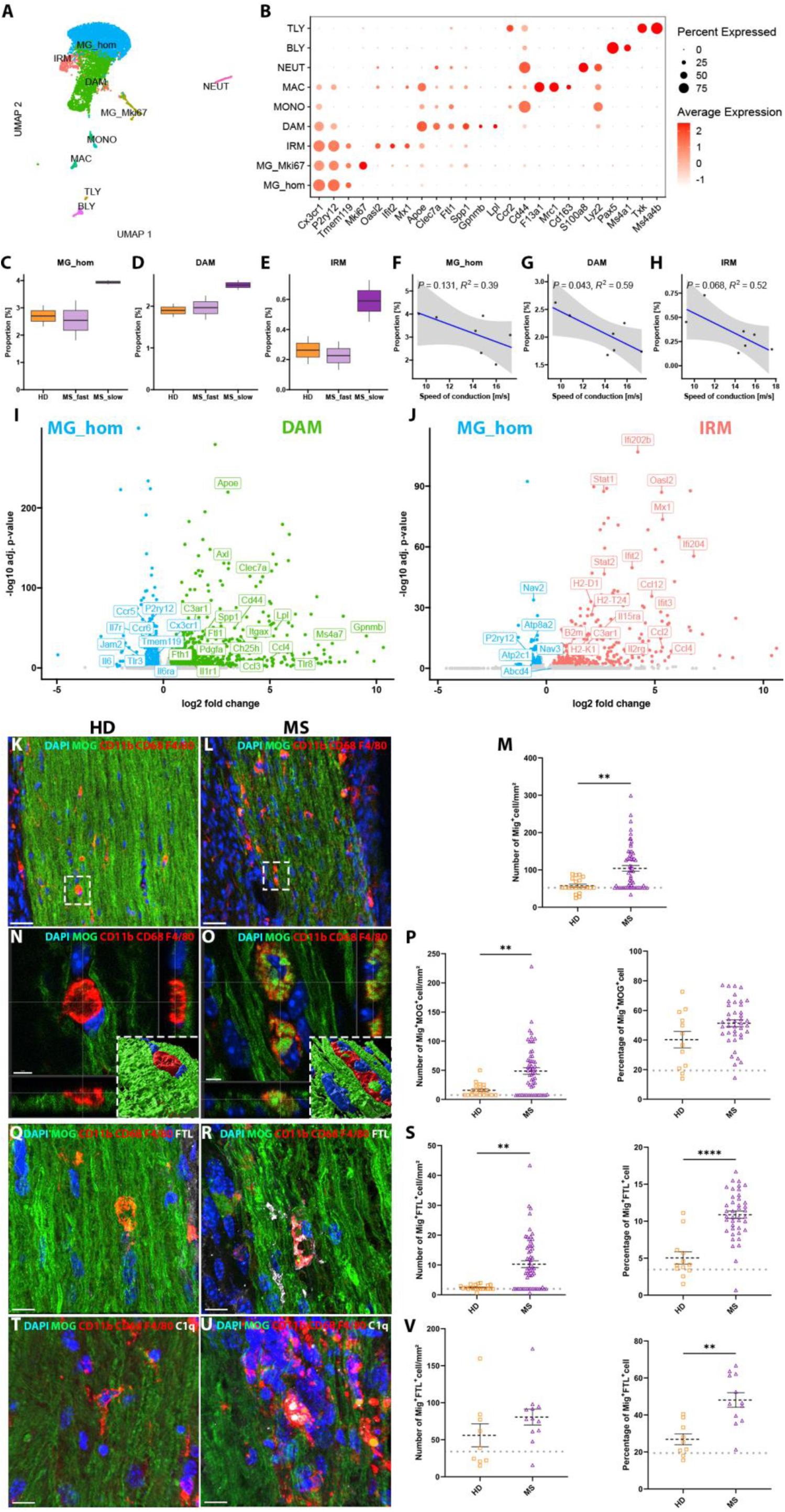
Emergence of disease associate microglia at the lesion rim after MS LY graft. **(A)** UMAP of microglia and immune cells after sub-clustering (5,797 nuclei). **(B)** Averaged gene expression of selected marker genes per cell population. **(C-E)** Boxplots highlighting relative cell abundance of homeostatic microglia (MG_hom), damage-associated microglia (DAM), and interferon-responsive microglia (IRM) signatures. **(F-H)** Cell abundance regressed against the speed of conduction. Strength and significance of associations were reported using the coefficient of determination (R²) and corresponding p-values. **(I-J)** Differential gene expression analysis showing selected genes enriched in (I) DAM (green) or (J) IRM associated microglia signatures (red) compared to homeostatic microglia (blue). **(K-V)** Immunostainings and quantifications of microglia and macrophages (CD11b, CD68, F4/80, red) at the lesion rim of HD LY (K, N, Q, T) and MS LY (L, O, R, U) grafted spinal cord lesions, 3 months post-surgery. Scale bars: 30µm (K-L), 5µm (N-O), and 10µm (Q-U). (K-P) Microglia showed continuous phagocytosis of myelin particles (MOG, green). (Q-S) Microglia accumulating ferritin light chain (FTL, white). (T-V) Microglia colocalization with complement factor C1q (white). **(M, P, S, V)** Quantifications of absolute cell abundance per mm^2^ (left panel) and percentage of myelin phagocytosing, FLT, or C1q positive signatures relative to total microglia (right panels) at the lesion rim. Each dot represents one mouse. Mean and SEM are displayed in black. The gray dotted line represents the averaged quantification of NG mice. Kruskal-Wallis test: ** p<0.01, **** p<0.0001.

To validate the overrepresentation of microglia as observed in snRNA-seq, we performed immunohistochemistry on myeloid cell markers (F4/80, CD11b, CD68, red Figure 5K-U), which confirmed an increased microglial density at lesion borders in MS LY grafted mice compared to NG and HD controls (Figure 5K–M). Myelin-containing myeloid cells (MOG green; Figure 5N–P) were also significantly more abundant in MS grafts, although their proportion relative to total myeloid cells was unchanged. Electron microscopy further captured microglia actively engulfing myelin at multiple stages, corroborating ongoing demyelination (Supplementary Figure 7A-C). Consistent with our snRNA-seq data showing overexpression of Ftl and C1qa in microglial subtypes, IHC confirmed increased representation of FTL+ and C1q+ microglia at lesion borders (Figure 5Q–V). Together, these results demonstrate that the lesion environment of MS LY grafted mice is enriched in activated microglia expressing FTL and C1q, which likely contribute to ongoing demyelination.

### Evidence of myelin pathology and impaired oligodendrocyte maturation in MS-grafted mice

Subclustering of the oligodendroglial lineage identified eight subpopulations, from OPCs to committed (COP), myelin-forming (MFOL), and mature oligodendrocytes (OL) (Figure 6A–B). Immature cell signatures, including OPCs and MFOLs, were enriched in MS samples associated to slow conduction speed. Their abundance negatively correlated with the SSEP conduction speed, while mature populations showed no such association (Figure 6C–F), suggesting that oligodendroglial immaturity leads to impaired nerve conduction.

**Figure 6:**
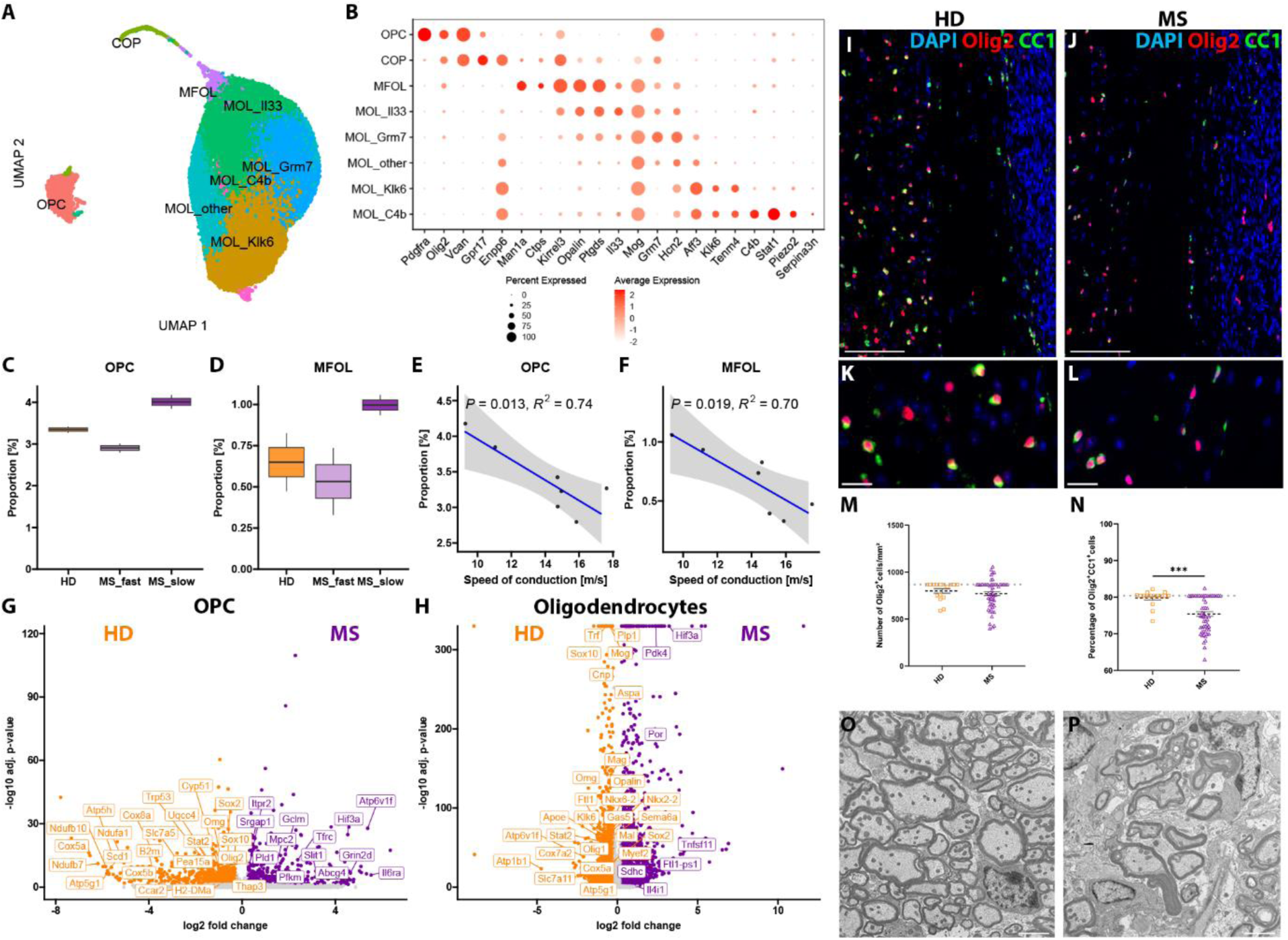
Myelin pathology and impaired oligodendrocyte maturation in MS-grafted mice. **(A)** UMAP of the OPC and oligodendrocytes after sub-clustering (47,319 nuclei). **(B)** Averaged expression of selected marker genes per cell population. **(C-D)** Boxplots highlighting relative cell abundance of (C) OPCs and (D) early myelin forming oligodendrocytes (MFOL) and their respective regression against the speed of conduction **(E-F)**. Strength and significance of associations were reported using the coefficient of determination (R²) and corresponding p-values. **(G-H)** Differential gene expression analysis of (G) OPC and (H) total oligodendrocytes, highlighting dysregulation of selected genes in MS (purple) compared to HD (orange). **(I-L)** Immunostaining of OLIG2 (red) and CC1 (green), HD LY grafted lesion (left panel) and MS LY grafted lesion (right panel). (I-J) View on global lesion (scale bar=100µm), (K-L) Zoom towards the border of the lesion (scale bar=20µm). **(M)** Mean number of Olig2 positive cells per mm² showed no difference in oligodendroglia cell abundance at the lesion border between MS and HD LY grafted lesions. **(N)** Percentage of Olig2^+^CC1^+^ cells highlighting a loss of mature (CC1 positive) lineage signature at the lesion rim of MS LY grafted lesions. Each dot represents one mouse, mean and SEM are displayed in black. The gray dotted line represents the averaged quantification of NG mice. Kruskal-Wallis test: ***: p<0,001. **(O-P)** Electron microscopy image at the lesion rim, highlighting disrupted compact myelin in MS LY grafted lesions (P) when compared to HD conditions (O). Scale bar = 2.5µm.

Differential gene expression analysis revealed downregulation of electron transport chain components (Cox-, Atp6v-, Nduf-, Uqcc-family of genes) and myelination-associated transcription factors (Olig2, Sox10) across OPCs and differentiated oligodendrocytes (Figure 6G-H). Mature oligodendrocytes additionally exhibited reduced expression of key genes involved in myelination (Cnp, Mog, Omg, Opalin, Mag, Klk6) and downregulation of cholesterol biosynthesis, consistent with diminished myelin maintenance. Immunostaining confirmed a lower abundance of mature Olig2+CC1+ cells at the lesion border of MS LY grafted mice despite comparable Olig2+ oligodendroglia number (Olig2 red, CC1 green, Figure 6I-N). Electron microscopy revealed disrupted myelin ultrastructure with myelin decompaction and swelling under MS conditions compared to HD (Figure 6O–P).

### Assessing the impact of individual donor LY on graft outcomes reveals patient-specific variability

Given that grafts of individual donor LY yielded variable outcomes, we applied a multiblock analysis incorporating behavioral, electrophysiological, and histological measurements (Figure 7) to visualize these effects. MCOA was used to visualize each mouse within a consensus space that integrates all modalities. This consensus space revealed that mice grafted with MS-derived LY segregated from those receiving HD LY on the first component, capturing 40% of the total variance (Figure 7A). Grouping mice by donor and calculating the cluster barycenters highlighted donor-specific effects: while some MS patients’ LY closely resembled HD profiles (e.g. MS9), others displayed a gradient of proximity to HD (Figure 7B), demonstrating that individual MS donors exert distinct effects on CNS. Calculating bootstrap confidence intervals confirmed the robustness of the metrics explaining the first MCOA component (Figure 7C). Specifically, the number of errors committed on the notched beam, as well as the abundance of myelin- or FTL-positive microglia at the lesion border, were strong predictors of MS LY effects. In contrast, faster speed of conduction, higher abundance of mature Olig2+CC1+ at the lesion border, and greater forelimb strength predicted HD LY graft outcomes.

**Figure 7.**
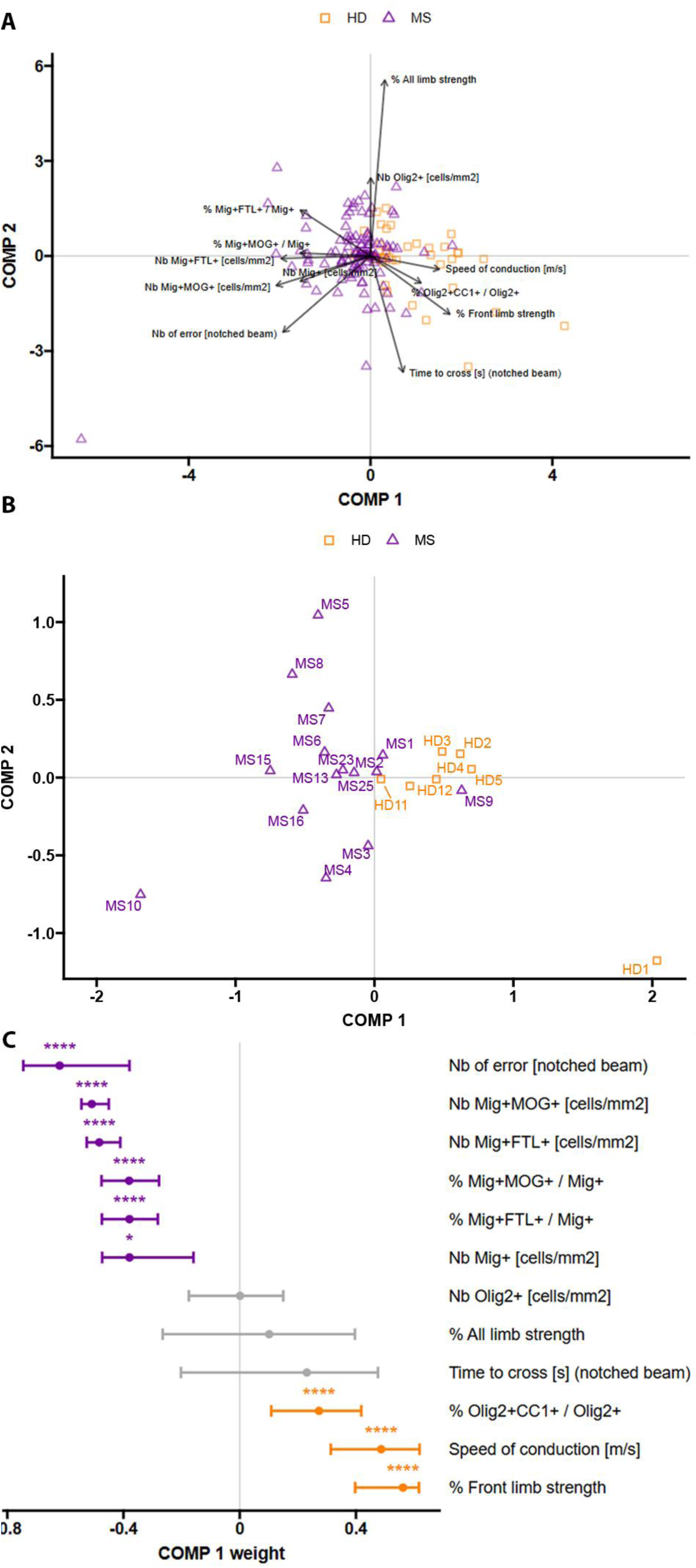
Heterogeneous effects of MS patient–derived lymphocyte grafts. **(A)** Biplot representation of the MCOA-based consensus space. Arrows highlight the experimental readouts that were assessed throughout this study. **(B)** Donor-specific barycenters computed from the consensus space. The experimental conditions MS and HD separated on component 1 (COMP 1) accounting for 40% total variation. **(C)** Variable contributions to the first component of the consensus space with bootstrap confidence intervals (weight); significant contributions are highlighted with stars *: p<0,05; **: p<0,01; ****: p<0,0001.

Together, these findings demonstrate that this chimeric graft model faithfully captures interpatient variability, providing a robust platform to dissect how individual MS donor LY differentially influence CNS pathology and function.

## Discussion

Animal models of MS provide distinct opportunities to explore different aspects of the disease. The EAE model has been invaluable for studying inflammatory processes in lesion formation, although the random distribution and timing of lesions complicate the study of remyelination (Procaccini et al., 2015). In contrast, the LPC model (Blakemore & Franklin, 2008; Hall, 1972) has been widely used to investigate remyelination in a minimally inflammatory setting. Building on these approaches, we combined focal demyelination with an inflammatory component by grafting human LY into LPC-induced lesions (El Behi*, Sanson* et al., 2017). Remarkably, LY from some MS patients inhibited remyelination, while LY from healthy donors (HD) or other MS patients did not interfere with this process. This highlighted donor-specific factors influencing repair and raised new questions about the long-term evolution of demyelinated lesions.

Three months after grafting, this model recapitulates several hallmark features of smoldering MS lesions. We observed lesions resembling mixed active/inactive MS pathology, with a glial-scared, inflammatory inactive center surrounded by an active rim (Kuhlmann et al., 2017). The active border contained iron- and myelin-laden myeloid cells, and LY from MS patients localized to the perivascular space—a characteristic feature of smoldering lesions (Absinta et al., 2021; Machado-Santos et al., 2018; Fransen et al., 2020; Frieser et al., 2022). In contrast, LY from HD rapidly disappeared, underscoring the requirement for both a permissive lesion environment and disease-specific lymphocyte traits to sustain their long-term retention.

The presence of LY perivascular cuffs in our model mirrors clinically relevant features of MS. In patients, such cuffs—composed of T and B cells differentiating into tissue-resident memory cells (T_RM_)—are associated with poorer outcomes (Hsiao et al., 2021; Machado-Santos et al., 2018; Magliozzi et al., 2023; Pignata et al., 2025; Smolders et al., 2018). These T_RM_ secrete factors that activate microglia and astrocytes, sustaining a chronic inflammatory environment that exacerbates tissue damage and impedes repair (Pignata et al., 2025; Woo et al., 2024). In line with these observations, MS LY in our model not only disrupted remyelination but also drove lesion expansion between three weeks and three months. While macrophages predominate during the initial phase of demyelination, resident microglia progressively take over, maintaining smoldering inflammation.

This progressive shift underscores the central role of microglia in sustaining chronic lesion activity. By three months, a distinct subset of activated microglia are present within the lesion rim environment. Marked by myelin phagocytosis, iron accumulation, and a shift from homeostatic to pro-inflammatory phenotypes, these cells closely resemble perilesional microglial cells that were described and detected by paramagnetic MRI signals in human smoldering lesions (Absinta et al., 2021; Klotz et al., 2023). Complement activation, including C1q expression, further supports the parallels between this model and human MS pathology (Absinta et al., 2021; Breij et al., 2008; Ingram et al., 2014; Lucchinetti et al., 2000; Prineas et al., 2001).

Within this environment, oligodendroglial lineage cells show clear signs of dysfunction. OPCs downregulate transcription factors essential for lineage progression, while mature oligodendrocytes reduce expression of myelin-related transcripts. These molecular alterations are consistent with prior observations reporting myelin abnormalities in normal appearing white matter ranging from compositional changes (Shaharabani et al., 2016; Teo et al., 2021) to structural defects, including myelin decompaction or swelling (Luchicchi et al., 2021, 2024)— features that we also identified at the rim of the lesion in our model. Moreover, we identified perturbations in stress-response pathways, including dysregulation of Hif3a, Hsp family members, and mitochondrial dysfunction, pointing to a state of chronic cellular stress. These findings are consistent with *post-mortem* studies demonstrating that oligodendrocytes in MS tissue persist under sustained stress conditions (Jäkel et al., 2019; Macnair et al., 2025; Pernin et al., 2024).

Finally, our model captures variations in donor LY effects, which may reflect some of the heterogeneity observed in patient profiles. Mice grafted with MS LY segregated into “slow” and “fast” SSEP subgroups, echoing inter-individual variability in remyelination efficiency that was observed clinically (Bodini et al., 2016). Recent transcriptomic studies further support this view: single-nucleus and bulk RNA sequencing have revealed that patient-specific transcriptional signatures, rather than lesion-specific ones, largely determine oligodendrocyte stress, maturation states, and glial interactions (Chen et al., 2025; Macnair et al., 2025). For example, one study (Macnair et al., 2025) stratified patients into four subgroups based on glial gene expression, uncovering donor-driven rather than lesion-specific patterns of oligodendrocyte stress and maturation, along with a continuum of glial transcriptional changes linked to microglial activation states Chen et al (Chen et al., 2025). classified MS donors by remyelination efficiency, identifying efficient versus poor remyelinating groups and pointing to actionable genes, particularly those involved in microglia–OPC/oligodendrocyte interactions, as therapeutic targets. Together, these findings highlight the dominant role of individual patient effects in shaping pathology and underscore the critical interplay between immune cells and oligodendrocytes.

In summary, our model provides a powerful experimental framework to investigate the transition from peripheral cell implication to chronic, compartmentalized CNS inflammation— a core driver of PIRA. By combining behavioral and physiological readouts with molecular and spatial transcriptomic analyses, it will enable non-invasive and longitudinal monitoring of lesion dynamics, offering unique opportunities to dissect the mechanisms underlying smoldering lesion formation. Beyond advancing our understanding of progressive inflammation, the model is well suited for testing therapeutic strategies targeting chronic lesion activity. Critically, by capturing patient-specific cellular interactions, it offers insights into the sources of disease heterogeneity and paves the way toward personalized therapeutic approaches in progressive MS.

## Limitation of the study

The clearing method and three-dimensional imaging enable us to detect human LY three months after the graft in MS LY grafted mice. However, the clearing method does not allow the use of all antibodies due to high tissue processing (such as high permeabilization), in addition, with few LY still present, we were not able to characterize them on classical immunohistochemistry experiments. Due to this limitation, we cannot conclude which subtypes of LY are the drivers of the development of smoldering lesions. To assess which subtypes of LY are necessary for the development of these lesions we could graft different subtypes of LY from one MS patient that we propose to induce smoldering lesions. However, we hypothesize that these LY are T_RM_ cells since they are still present in the tissue even three months after the graft. In our model, LY were found at the perivascular space, similar to T_RM_ that were localized to perivascular spaces of activated and smoldering lesions and that were implicated in the compartmentalized inflammation.

## Acknowledgments

We are grateful to all the patients and volunteers that participated in this study, and to the members of the Zujovic/Nait Oumesmar and Zujovic /Mochel team for their support. We are indebted to the Bouvet-Labruyère family for their constant support to MS research at the Paris Brain Institute. The authors wish to thank the cell culture (CELIS), sequencing (i-GENSEQ), Data analysis Core (DAC, RRID:SCR_026138), animal (PhenoParc), imaging (ICM-Quant, RRID:SCR_026393), electrophysiology (e-PHYS) facilities but also the fundraising, scientific affairs and the administrative department of the Paris Brain Institute. We want to thank also Sarah Taieb Tamacha and Alexander Balcerac from the CIC Neurosciences for their efficient management of MS patient recruitment of the MS-BIOPROGRESS cohort and for supervision of blood sampling from MS patients.

## Author contributions

VZ designed and supervised the project. OP performed data acquisition, analysis, and drafted and wrote the manuscript. VZ, MN wrote, reviewed and edited the manuscript. CL supervised the MSBIOPROGRESS cohort management and patient recruitment. CB, DA, NS, AL and MN participated in data acquisition and analysis. AT supervised the statistical analysis. All authors discussed the results and commented on the manuscript.

## Abbreviation

CNS: Central Nervous System
COP: Committed oligodendrocyte precursor cell
DAM: Disease associated microglia
DEG: Differential gene expression
EAE: Experimental Autoimmune Encephalomyelitis
EM: Electron microscopy
HD: Healthy donor
IHC: Immunohistochemistry
IRM: Interferon responsive microglia
LPC: Lysophosphatidylcholine
LY: Lymphocytes
MCOA: Multiple Co-inertia analysis
MFOL: Myelin forming oligodendrocyte
MS: Multiple Sclerosis
NG: Non-grafted
OL: Oligodendrocytes
OPC: Oligodendrocytes Precursors Cell
PBMC: Peripheral Blood Mononuclear cell
PIRA: Progression Independent of Relapse Activity
PPMS: Primary Progressive Multiple Sclerosis
RAW: Relapse Associated Worsening
RGCCA: Regularized generalized canonical correlation analysis
RRMS: Relapsing Remitting Multiple Sclerosis
SnRNAseq: Single nuclei RNA sequencing
SPMS: Secondary Progressive Multiple Sclerosis
SSEP: Somatosensory evoked potential.

