## Supplementary material for "Patient-derived lymphocytes drive smoldering lesion pathology in a chimeric multiple sclerosis mouse model"

### Supplementary Figures :

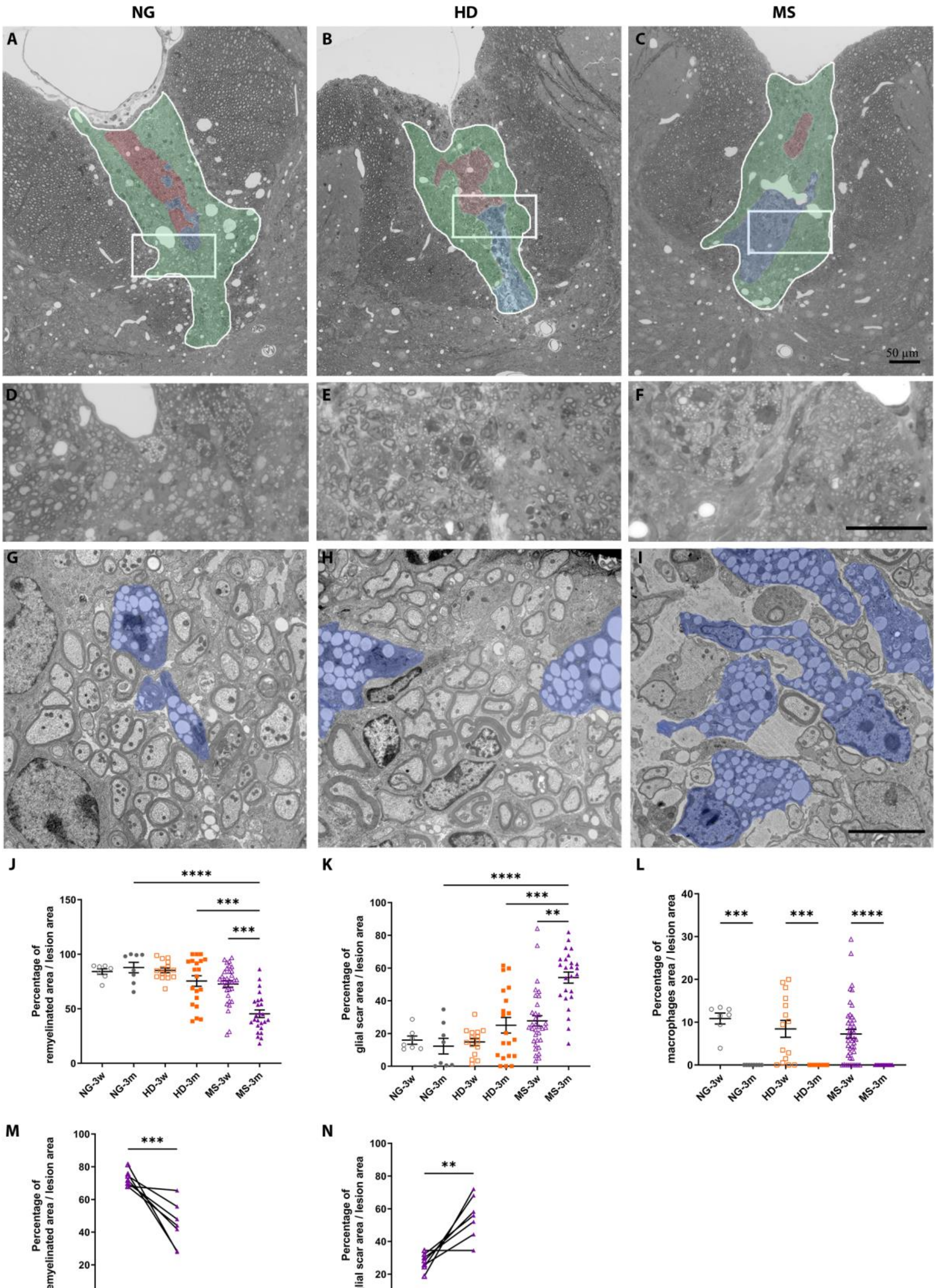

***Supplementary Figure 1: Lesion progression between 3 weeks and 3 months.* *(A-C)*** *Semi thin sections of the dorsal horn of spinal cords lesions from NG mice, (A, D, G; n= 7 mice), and after grafting HD LY (B, E, H; n= 5 HD donors, 2-3 mice per donor); and MS patient LY (n= 7 MS donors, 2-4 mice per donor) 3-weeks post-surgery. Highlighted are the remyelinated area by oligodendrocytes (green) or Schwann cells (red), as well as macrophages (blue).* ***(D-F)*** *Zoom inside the lesion indicated by the square above.* ***(G-I)*** *Macrophages with vacuoles (blue) at the lesion rim.* ***(J)*** *Mean percentage of remyelinated area in reference to total lesion area at 3-weeks and 3-months post-surgery per condition.* ***(K)*** *Mean percentage of glial scar area in reference to total lesion area at 3-weeks and 3-months post-surgery per condition.* ***(L)*** *Mean percentages of macrophage area in reference to total lesion area at 3-weeks and 3-months of each group: NG (n= 7 mice at 3 weeks and 3 months, gray), HD (n= 15 mice at 3 weeks and 20 mice at 3 months, orange) and MS (n= 33 mice at 3 weeks and 25 mice at 3 months, purple). One way ANOVA followed by Tukey's post-hoc: ***: p<0,001; ****: p<0,0001.* ***(M)*** *Loss of remyelinated area and* ***(N)*** *gain in glial scar area in MS LY grafted mice 3-months post-surgery compared to 3-weeks post-surgery. Mice that were grafted with LY from the same donor were averaged. Wilcoxon test: **: p<0,01; ***: p<0,001.*

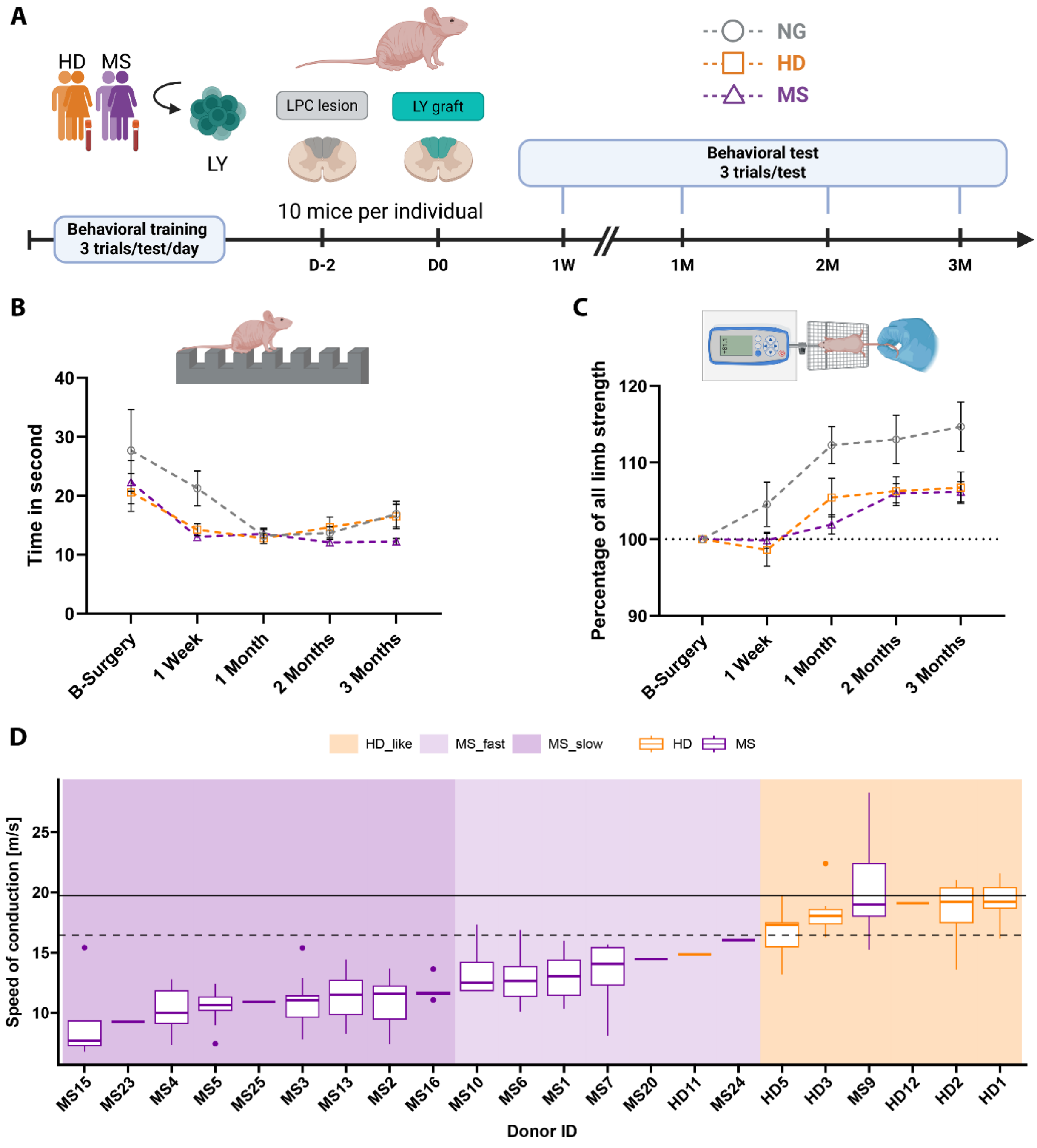


***Supplementary Figure 2: Behavioral and physiological assessment. (A)*** *Experimental setup of behavioral assessments (NG gray, HD orange, MS purple). For each behavioral test, three trials per time point and mouse were assessed and averaged.* ***(B)*** *Mean time to cross the notched beam in second.* ***(C)*** *Relative all limb strength in percentage to baseline assessment.* ***(D)*** *Boxplots highlighting the SSEP conduction velocity through the spinal cord lesion. Mice were grouped into the LY donors to infer donor specific effects on lesion severity. Donors were order according to median speed of conduction, highlighting MS donors that induced severe lesions associated to slow conduction velocity (< 12 m/s, MS_slow, dark purple), MS donors associated to faster conduction velocity (12-17 m/s, MS_fast, light purple), and donors associated to conduction velocities resembling healthy remyelination outcomes (> 17 m/s, HD-like, orange). The dotted line represents median conduction velocity of NG mice (16.4 m/s), the solid line represents median conduction velocity of non-lesioned mice (19.75 m/s).*

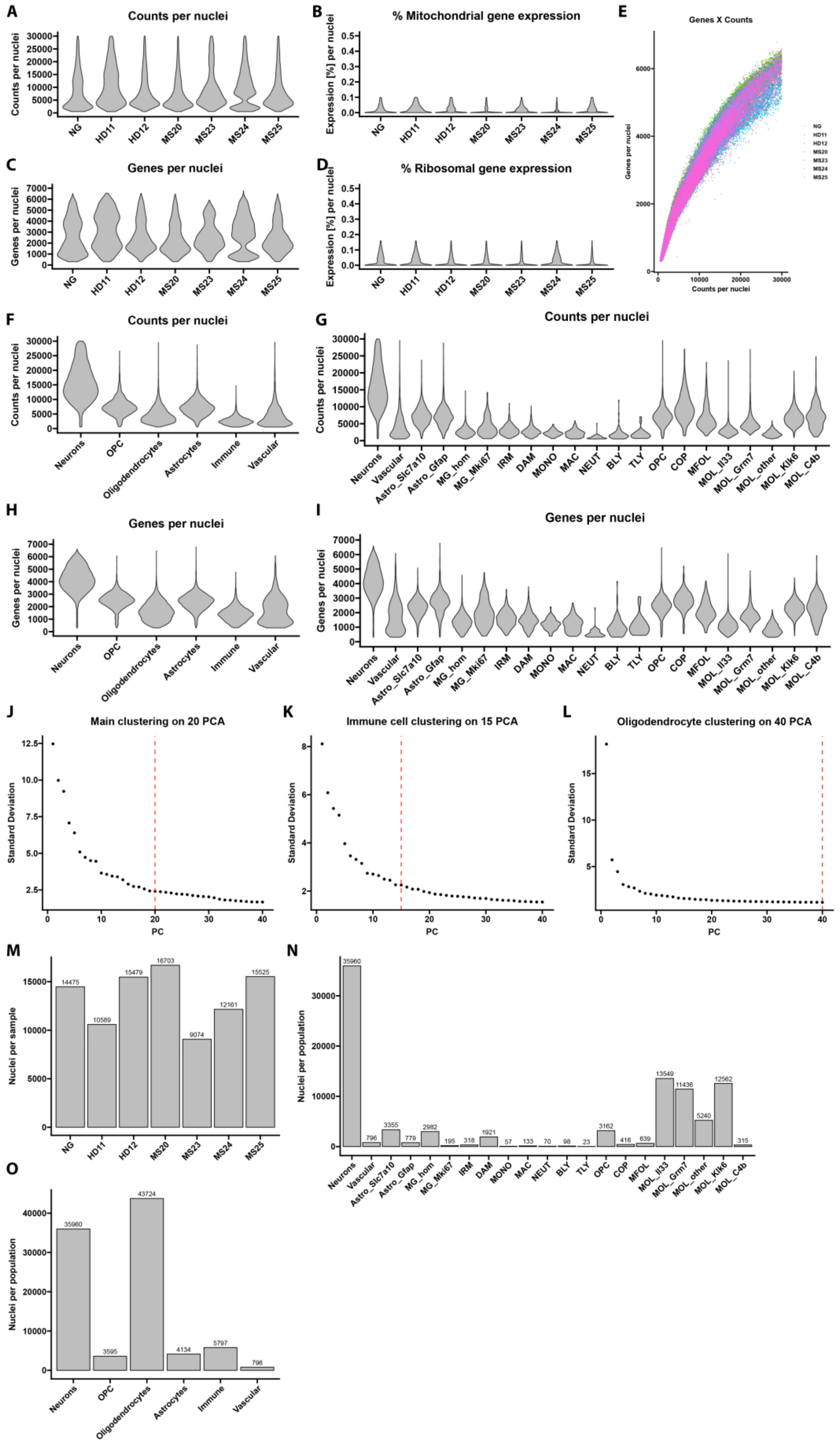


***Supplementary Figure 3: Quality control of the single nuclei RNA seq dataset. (A, F, G)*** *Number of sequenced UMI counts per nuclei, split per sample (A), broad CNS annotated cells (F), and sub-populations (G). Nuclei were*

*filtered for min 500 and max 30 000 UMI counts.* ***(C, H, I)*** *Number of genes expressed per nuclei, split per sample (C), broad CNS annotated cells (H), and sub-populations (I). Nuclei were filtered for min 300 and max 7 000 genes expressed. (****B****) Percentage of mitochondrial genes expressed per nuclei and split per sample. A threshold of 0.1% was chosen for nuclei filtering. (****D****) Percentage of ribosomal genes expressed per nuclei and split per sample. A threshold of 0.16 % was chosen for nuclei filtering.* ***(E)*** *Scatter plot highlighting the number of genes expressed per nuclei and the numbers of UMI counts sequenced per nuclei and split by the samples.* ***(J, K, L)*** *Elbow plots highlighting the number of PCA chosen for downstream clustering of the global dataset (J), as well as the sub-clustering of immune cells (K) and oligodendroglia (L).* ***(M, N, O)*** *Number of nuclei per clustered CNS population (M), sub-populations (N), and samples (O).*

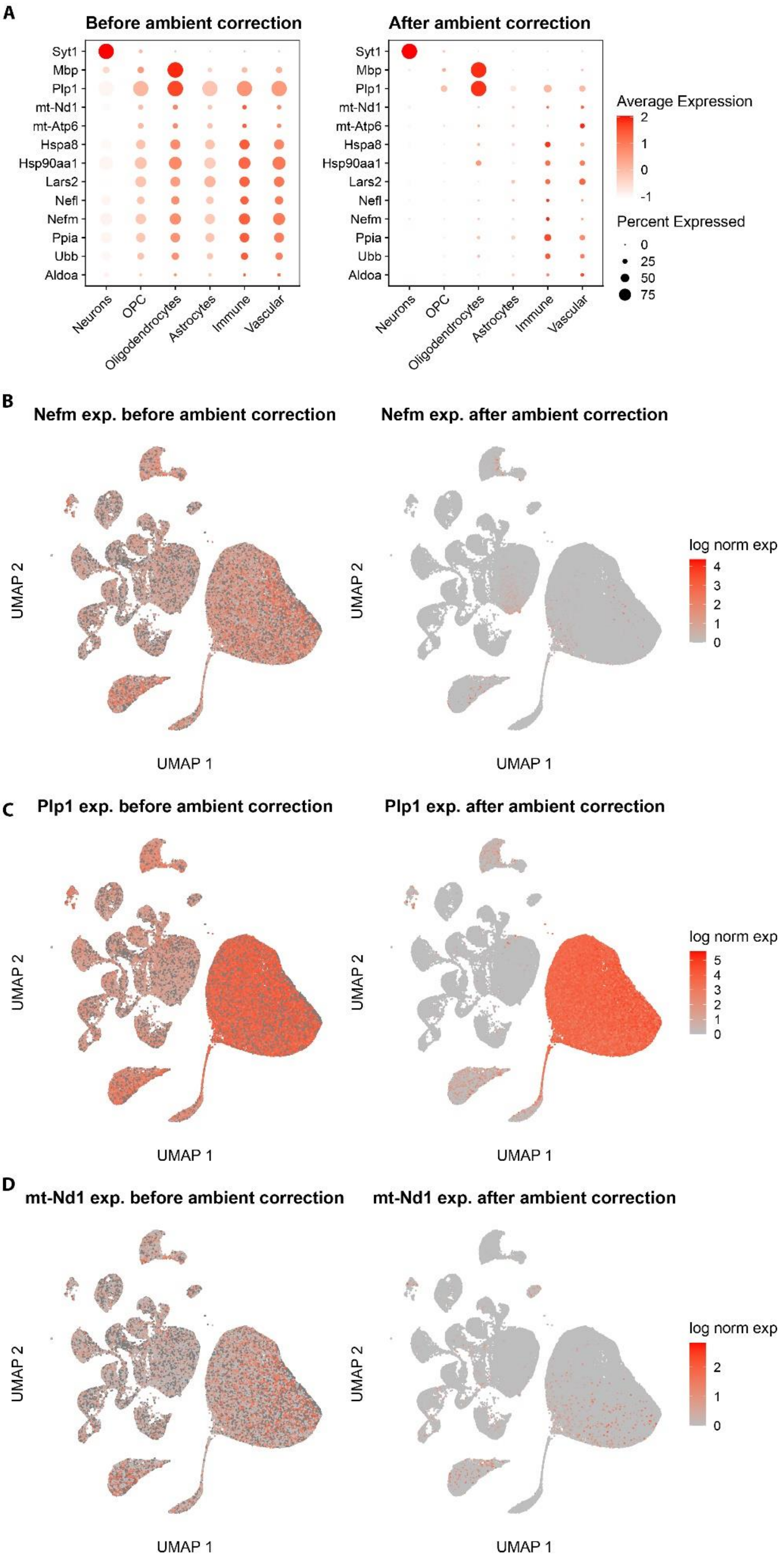


***Supplementary Figure 4: Quality control of ambient RNA expression before and after cellbender background correction. (A)*** *Dot plots highlighting averaged expression levels of selected genes that were found highly expressed in empty droplets, mapped to the annotated clusters before (left) and after (right) ambient RNA*

*correction.* ***(B-D)*** *Log normalized gene expression of Nefm (B), Plp1 (C), and mt-Nd1 (D) per single nuclei mapped to the UMAP embedding before (left) and after (right) ambient RNA correction.*

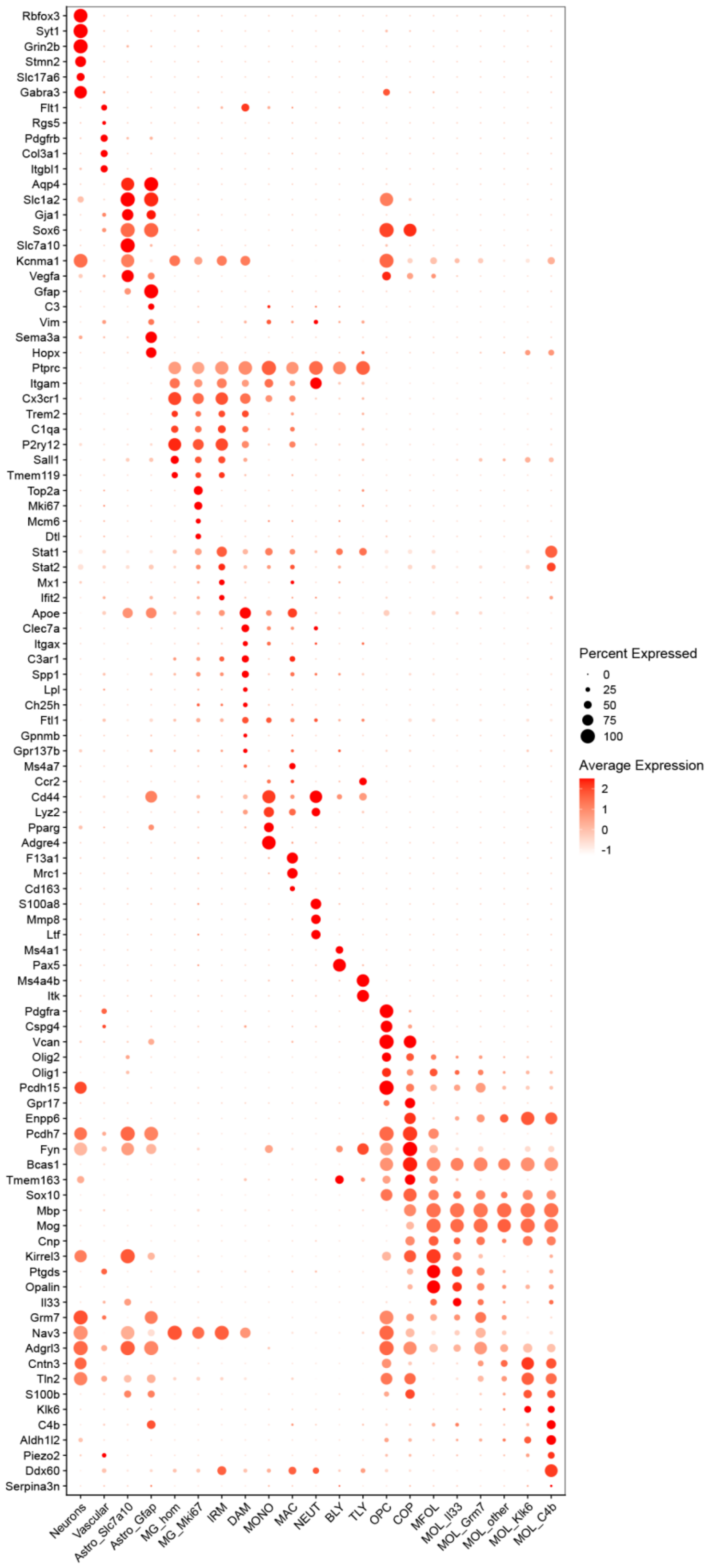
Rbfox3
Syt1
Grin2b
Stmn2
Slc17a6
Gabra3
Flt1
Rgs5
Pdgfrb
Col3a1
Itgbl1
Aqp4
Slc1a2
Gja1
Sox6
Slc7a10
Kcnma1
Vegfa
Gfap
C3
Vim
Sema3a
Hopx
Ptprc
Itgam
Cx3cr1
Trem2
C1qa
P2ry12
Sall1
Tmem119
Top2a
Mki67
Mcm6
Dtl
Stat1
Stat2
Mx1
Ifit2
Apoe
Clec7a
Itgax
C3ar1
Spp1
Lpl
Ch25h
Ftl1
Gpnmb
Gpr137b
Ms4a7
Ccr2
Cd44
Lyz2
Pparg
Adgre4
F13a1
Mrc1
Cd163
S100a8
Mmp8
Ltf
Ms4a1
Pax5
Ms4a4b
Itk
Pdgfra
Cspg4
Vcan
Olig2
Olig1
Pcdh15
Gpr17
Enpp6
Pcdh7
Fyn
Bcas1
Tmem163
Sox10
Mbp
Mog
Cnp
Kirrel3
Ptgds
Opalin
Il33
Grm7
Nav3
Adgrl3
Cntn3
Tln2
S100b
Klk6
C4b
Aldh1l2
Piezo2
Ddx60
Serpina3n
Neurons
Vascular
Astro_Slc7a10
Astro_Gfap
MG_hom
MG_Mki67
IRM
DAM
MONO
MAC
NEUT
BLY
TLY
OPC
COP
MFOL
MOL_Il33
MOL_Grm7
MOL_other
MOL_Klk6
MOL_C4b
Percent Expressed
0
25
50
75
100
Average Expression
2
1
0
-1

***Supplementary Figure 5: Marker expression of annotated clusters.*** *Dot plot highlighting averaged gene expression of selected genes used for cluster annotation and differentiation.*

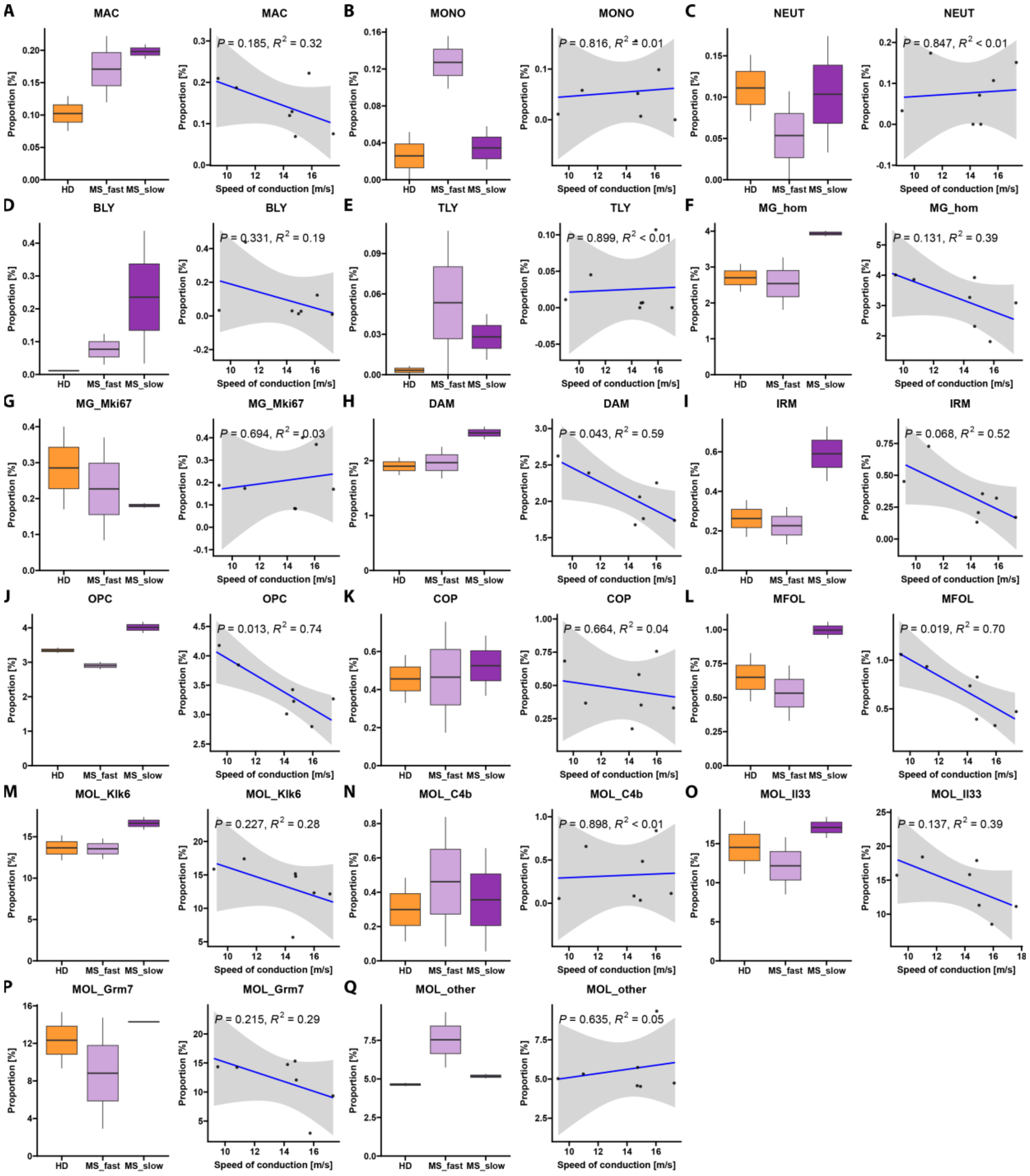


***Supplementary Figure 6: Cell abundance and SSEP correlations of subclustered cell types.*** *Boxplots highlighting relative cell abundance in percent and their respective correlation with the SSEP inferred speed of conduction of (A) Macrophages (Mac), (B) Monocytes (MONO), (C) Neutrophils (NEUT), (D) B lymphocytes (BLY), (E) T lymphocytes (TLY), (F) homeostatic microglia (MG_hom), (G) proliferating microglia (MG_Mki67), (H) damage-associated microglia (DAM), (I) Interferon-responsive microglia (IRM), (J) oligodendrocyte precursor cells (OPC), (K) committed oligodendrocyte precursor (COP), (L) early myelin forming oligodendrocytes (MFOL), and mature oligodendrocytes (OL) (M-Q). Regression strength and significance of associations were reported using the coefficient of determination ($R^2$) and corresponding p-values.*

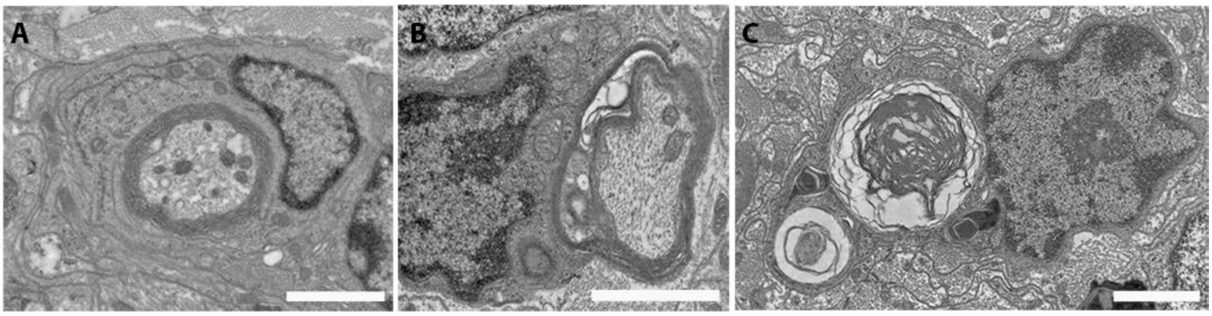


***Supplementary Figure 7: Ultrastructural features of myelin phagocytosis by microglial cells. (A-C)*** *Ultra-thin sections of the dorsal horn of spinal cord of Nude mice after the lesion and grafting of MS patient LY 3 months after the surgery. Each image illustrates a distinct phase of myelin phagocytosis: (A) the engulfment of a myelinated axon, (B) the unwrapping of myelin, and (C) the process of myelin phagocytosis.*

### Supplementary tables:

***Supplementary table 1: Individual description of multiple sclerosis patient's cohort.***

| Code | Disease_Type | Sex | Age | Treatment |
|---|---|---|---|---|
| MS1 | RR | F | 43 | None |
| MS2 | RR | F | 32 | None |
| MS3 | RR | F | 51 | None |
| MS4 | RR | F | 29 | None |
| MS5 | RR | F | 70 | None |
| MS6 | RR | F | 69 | None |
| MS7 | RR | M | 46 | Glatiramer acetate |
| MS8 | RR | F | 36 | Dimethyl fumarate |
| MS9 | PP | F | 50 | None |
| MS10 | RR | F | 23 | Dimethyl fumarate |
| MS13 | RR | M | 44 | None |
| MS15 | RR | F | 33 | Dimethyl fumarate |
| MS16 | RR | F | 49 | Glatiramer acetate |
| MS20 | RR | F | 41 | None |
| MS23 | RR | F | 54 | Dimethyl fumarate |
| MS25 | RR | M | 31 | None |
| MS24 | RR | F | 47 | Diroximel fumarate |
| HD1 | \ | F | 34 | \ |
| HD2 | \ | F | 51 | \ |
| HD3 | \ | F | 49 | \ |
| HD4 | \ | F | 39 | \ |
| HD5 | \ | F | 25 | \ |
| HD11 | \ | M | 29 | \ |
| HD12 | \ | F | 35 | \ |

*Supplementary table 2: Antibodies used in the study.*

| Antibody | Dilution | Target | Isotype | Blocking/Permeabilisation solution | Source (Reference) |
|---|---|---|---|---|---|
| C1q | 1/50 | Complement | Mouse IgG2b-Biotin | PBS/BSA 4% Titron 0,2% | Thermofischer (MA1-40312) |
| CC1 | 1/100 | Mature OL | Mouse IgG2b | PBS/BSA 4% Titron 0,2% | Millipore (OP80) |
| CD11b | 1/400 | Macrophages/ Microglia | Rat IgG | PBS/BSA 4% Titron 0,2% | Bio-rad (MCA74G) |
| CD69 | 1/400 | Macrophages/ Microglia | Rat IgG | PBS/BSA 4% Titron 0,2% | Bio-rad (MCA1957) |
| Coll IV | 1/400 IHC<br>1/200 Clearing | Blood vessels | Rabbit IgG | PBS/BSA 4% Titron 0,2% | Abcam (ab19808) |
| F4/80 | 1/100 | Macrophages/ Microglia | Rat IgG | PBS/BSA 4% Titron 0,2% | Bio-rad (MCA497R) |
| FTL | 1/100 | Iron load | Rabbit IgG | PBS/BSA 4% Titron 0,2% | Abcam (ab69090) |
| HLA-DP,DQ, DR (Clone CR3/43) | 1/100 | Human LY | Mouse IgG1 | PBS/BSA 4% Titron 0,2% | Dako (M0775) |
| MOG | 1/100 | Myelin | Mouse IgG1 | PBS/BSA 4% Titron 0,2% | Home made |
| Olig2 | 1/200 | OL lineage | Rabbit IgG | PBS/BSA 4% Titron 0,2% + Antigen retrieval | Millipore (AB9610) |
